# Morphology of migration: Associations between wing, and bill morphology and migration in kingbirds (*Tyrannus*)

**DOI:** 10.1101/2021.04.06.438708

**Authors:** Maggie P. MacPherson, Alex E. Jahn, Nicholas A. Mason

## Abstract

Morphology is closely linked to locomotion and diet in animals. In animals that undertake long-distance migrations, limb-morphology is under selection to maximize mobility and minimize energy expenditure. Migratory behaviors also interact with diet, such that migratory animals tend to be dietary generalists, while sedentary taxa tend to be dietary specialists. Despite a hypothesized link between migration status and morphology, phylogenetic comparative studies have yielded conflicting findings. We tested for evolutionary associations between migratory status and limb and bill morphology across kingbirds, a pan-American genus of birds with migratory, partially migratory, and sedentary taxa. Migratory kingbirds had longer wings, in agreement with expectations if selection favors improved aerodynamics for long-distance migration. We also found an association between migratory status and bill shape, such that more migratory taxa had wider, deeper, and shorter bills compared to sedentary taxa. However, there was no difference in intraspecific morphological variation among migrants, partial migrants, and residents, suggesting that dietary specialization has evolved independently of migration strategy. The evolutionary links between migration, diet, and morphology in kingbirds uncovered here further strengthen ecomorphological associations that underlie long-distance seasonal movements in animals.

## Introduction

Animal movement is linked to morphology at various taxonomic scales. At a macroevolutionary scale, streamlined or aerodynamic body shapes have been associated with the evolution of long-distance migration in fish (Chapman *et al.*, 2015), insects (Johansson, Söderquist, & Bokma, 2009), and birds (Fiedler, 2005). Within species, migratory distance has also been associated with streamlined body shapes in fish (e.g., Crossin *et al.*, 2004) and aerodynamic shapes in birds (Voelker, 2001; Minias *et al.*, 2015; Vágási *et al.*, 2016). While certain taxa with long-distance movements exhibit strong selection for energy-efficient body shapes, this is not universal (Mulvihill & Chandler, 1990; Mönkkönen, 1995; Wang & Clarke, 2015). Thus, there is a persistent need to expand the taxonomic breadth of studies linking migration and morphology to better understand which lineages exhibit migration-morphology associations, and why these associations vary among taxa.

Morphology is also shaped by foraging strategies. For example, dietary niche is associated with head and body shape in fish (Knudsen *et al.*, 2011; Závorka *et al.*, 2020), birds (Felice *et al.*, 2019), and mammals (Swanson, Oliveros, & Esselstyn, 2019). Additionally, phenotypic plasticity is expected in dietary generalists, as shown in comparative common-garden experiments in stickleback minnows (Svanbäck & Schluter, 2012). In populations recently released from interspecific competition, such as island colonizers (Wilson, 1961; Clegg & Owens, 2002), phenotypic plasticity is thought to support increased morphological variation to limit intraspecific competition (i.e., ‘niche variation hypothesis’, Van Valen, 1965). Migratory lineages of dietary generalists may have strong preferences for food resources that are easiest to access (Sherry, 1984; Levey & Stiles, 1992; Bell, 2011) or that are superabundant (Moreau, 1952; Morse, 1971; Willis, 1974). In contrast, sedentary individuals may mitigate intra- and interspecific competition by seeking temporally stable food resources in microhabitats that are buffered from environmental fluctuation (e.g., temperature fluctuations; Bell, 2011). Comparisons across avian species (Levey & Stiles, 1992) and families (Chesser & Levey, 1998) have shown migratory taxa that forage on seasonally variable resources tend to exhibit more morphological variation, presumably associated with opportunistic foraging across a wider dietary breadth among or within individuals (Bell, 2011).

Testing how migration shapes morphological variation among taxa requires a phylogenetic comparative framework with comprehensive inter- and intraspecific sampling. Studies comparing migratory to sedentary birds support that long-distance migration favors longer (Rayner, 1988; Wiedenfeld, 1991; Pérez-Tris & Tellería, 2001; Förschler & Bairlein, 2011; Tellería *et al.*, 2013), and more pointed (Carvalho Provinciato, Araújo, & Jahn, 2018; Gómez-Bahamón *et al.*, 2020b) wings for increased aerodynamics. Hypotheses that link bill morphology and migratory status are less well-studied, but hinge on differences in diet that covary with migratory status (Bell, 2011; but see Cox, 1968; Herrera, 1978; Leisler, 1990). Sedentary insectivorous taxa often have longer bills, presumably to improve closing speed for capturing highly mobile prey (Leisler, 1990). In contrast, migratory taxa may have shorter bills for capturing slow-moving prey like caterpillars to feed young during the breeding season and small, abundant arthropods during winter (Thiollay, 1988; Leisler, 1990). Furthermore, studies on bill size in relation to foraging strategies disagree, such that both narrow and wide bills are associated with generalist diets (Tellería & Carbonell, 1999; Tellería *et al.*, 2013), and bill size differs between isolated specialist populations of the same species (Alonso *et al.*, 2020).

Early assessments of morphological variation among migratory and sedentary avian species did not account for their shared evolutionary history (Cox, 1968, 1985; Leisler, 1990). Boyle and Conway (2007) advanced this approach by performing phylogenetically independent contrasts to address selective pressures for the evolution of migration between species (see also Gómez *et al.*, 2016; Vágási *et al.*, 2016). However, the literature on the role of migratory status in shaping morphological phenotypes in birds draws predominantly from comparisons among distantly related taxa (e.g., Rappole’s 1995; critique of Herrera 1978). Thus, detailed comparisons among taxa that differ in migratory status within a modern phylogenetic comparative framework are necessary for a more comprehensive test of how migration is associated with morphology at different taxonomic scales (e.g., Bolnick *et al.*, 2007).

We used phylogenetic comparative methods to test for ecomorphological associations in wing and bill morphology among migratory, partially migratory, and sedentary kingbirds (*Tyrannus*; Fitzpatrick *et al.*, 2004). Kingbirds are flycatchers (Tyrannidae) with considerable variation in migration status and morphology within and among species, as well as a rich body of literature linking ecology and morphology (Fitzpatrick & Schauensee, 1980; Fitzpatrick, 1981; Sherry, 1984; Fitzpatrick, 1985; Cintra, 1997; Fitzpatrick *et al.*, 2004; Gabriel & Pizo, 2005; Carvalho Provinciato *et al.*, 2018; Gómez-Bahamón *et al.*, 2020b). As aerodynamic theory predicts that longer, more pointed wings and shorter tails are more efficient for long-distance migratory flights (Norberg, 1995; Pennycuick, 2008), we expected migratory taxa to have longer and more pointed wings compared to sedentary taxa (Kipp, 1942, 1958; Winkler & Leisler, 1992; Mönkkönen, 1995 and references therein; Lockwood, Swaddle, & Rayner, 1998). Tails of long-distance migrants should also be shorter than more sedentary individuals to reduce drag during long-distance flights (Rayner, 1988; Norberg, 1990; Winkler & Leisler, 1992; Förschler & Bairlein, 2011). However, tails are targets of sexual selection (Winquist & Lemon, 1994; Mobley, 2002), and tail lengths are often unassociated with migration strategy (e.g., Voelker, 2001; Neto *et al.*, 2013). Within-population variation in flight capability could be associated with migration (Fernández & Lank, 2007) or foraging ecology (Hromada & Tryjanowski, 2003), so we also tested whether more migratory taxa have more variable wing morphology compared to less migratory taxa.

Bill morphology is closely tied to foraging niche (Snow, 1953, 1954; Lack, 1971) and bill size and shape is typically related to broad dietary categories in birds (e.g., >50% insectivorous, granivorous, frugivorous etc.; e.g., Reaney *et al.*, 2020). We therefore predicted that there would be no relationship between migratory status and bill size because all kingbirds are primarily insectivorous. Additionally, migration may impart an ‘ecological release’ associated with the evolution of a more variable, opportunistic diet (Bell, 2011). We thus predicted that more migratory taxa will have higher coefficients of variation in bill morphology compared to less migratory taxa.

## Methods

### Phylogeny construction

We extracted the clade of 15 kingbird taxa from the Harvey et al. (2020) suboscine phylogeny to construct a kingbird phylogeny that includes 28 operational taxonomic units (OTUs). Harvey et al. (2020) included all species (13/13) but only 15% of the subspecific diversity (3/20). We therefore added *T. savana* taxa following phylogenetic relationships and branch lengths estimated by Gómez-Bahamón (2020a). The remaining subspecies were added as polytomies assuming a most recent common ancestor (MRCA) of 0.5 million years from members of their species group (Figure 1A). To test the sensitivity of analyses to this assumption, we tested a range of different dates for the MRCA for the added subspecies, which did not change the results (Supplementary Information 1).

**Figure 1.**
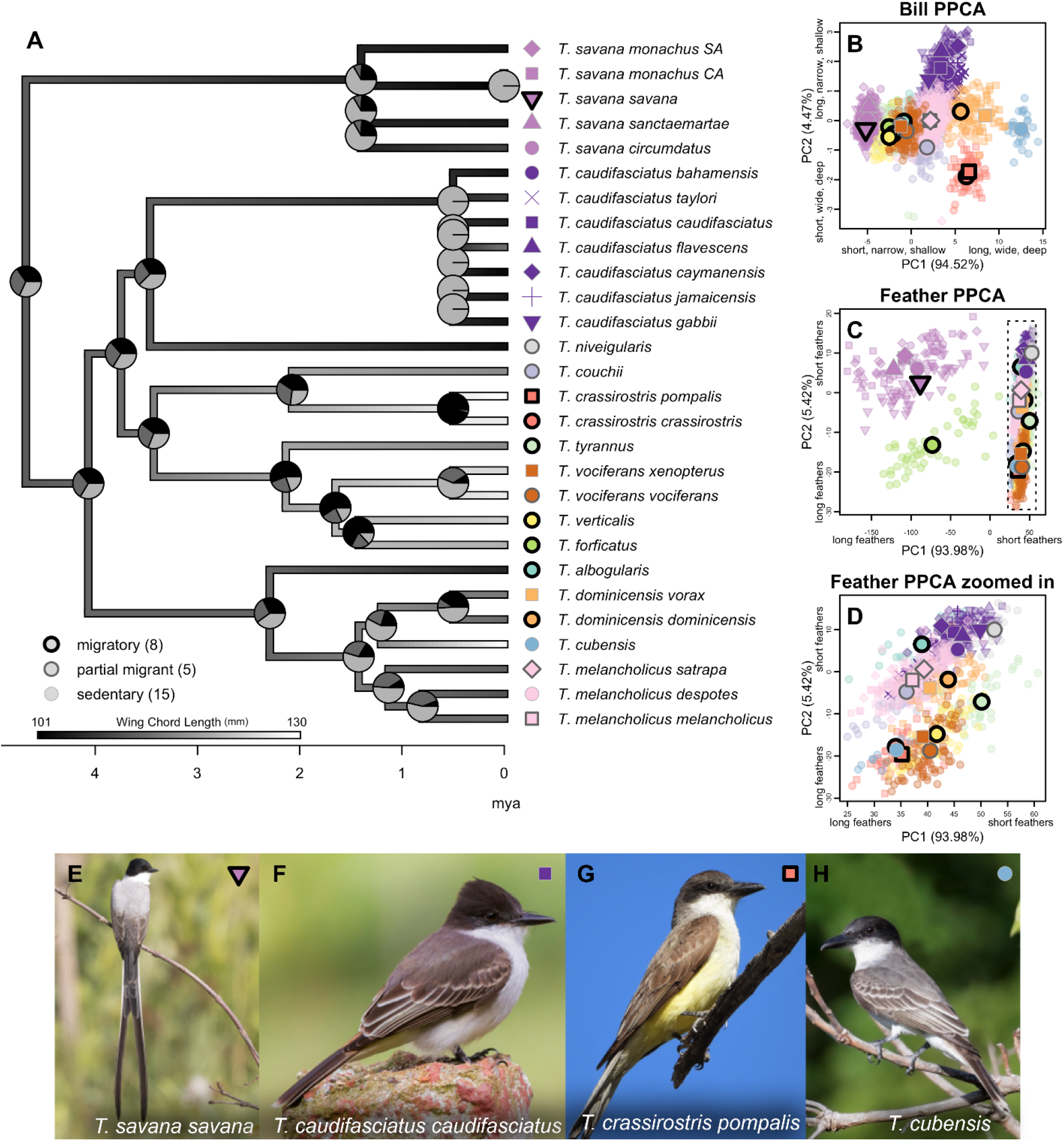
Ancestral state reconstruction of migration ecology category and wing chord length, as well as ordination of 28 kingbird OTUs for bill (length, width, depth) and feather (wing chord length, tail length, and Kipp’s distance). A) Kingbird phylogeny showing ancestral state reconstruction of migration ecology strategy (pie charts at each node), and wing chord length (branch greyscale). B-D) PPCA plots showing individuals color-coded to species (identified at the tip labels of the phylogeny), shape-coded to subspecies, and their status as migratory, partially migratory, or sedentary is distinguished via the shape outline. Pie charts for each node in the phylogeny show the ancestral state reconstruction of migration type. E-H) photos of representative kingbird taxa with corresponding color- and shape-coded points in the top right of each photo. Taxa are as follows: E) *Tyrannus savana savana* (photo credit: Rodrigo Conte), F) *Tyrannus caudifasciatus caudifasciatus* (photo credit: Yeray Seminario / Whitehawk), G) *Tyrannus crassirostris pompalis* (photo credit: Martin Molina), and H) *Tyrannus cubensis* (photo credit: Dubi Shapiro).

### Morphological measurements

We measured bill length, width, and depth at the distal end of the nares. We also measured unflattened wing chord length, Kipp’s distance (a measure of wing pointedness: the distance between the tip of the first secondary feather to the tip of the longest primary feather; Kipp, 1942, 1958; Baldwin *et al.*, 2010), tail length, and tarsus length on 2108 study skins from across the ranges of each species and subspecies (28 operational taxonomic units (OTUs); Figure 1A; Table S1.1). MM measured 2008 specimens, and identified all individuals to the lowest level of taxonomic identification (using Clements *et al.*, 2019) and classified each individual as migratory, partially migratory, or sedentary (*sensu* Fitzpatrick *et al.*, 2004). Partially migratory taxa are those in which individuals vary in migratory tendency (Boyle, 2008). JIGA (see Acknowledgements) measured 100 specimens to improve sampling of some taxa. Bill and tail measurements were taken twice and were confirmed to be within one mm of one another. Both right and left wing chords and tarsus lengths were measured and averaged. Measurements by JIGA were only taken once. We measured tail length as the longest rectrix to the nearest 0.1 cm (Pyle *et al.*, 1997). For most measurements, we used a Mitutoyo brand IP 67 digital calipers (part number 573-271) with a range of up to 15.24 cm, with 0.00127 cm resolution. For tails longer than 15.24 cm, we used a 30.48 cm stainless steel ruler placed between the two middle rectrices. When tails were longer than 30.48 cm photos were taken of the tails above a 0.64 x 0.64 cm square grid with the calipers measuring to their extent and ImageJ was used to calculate the full tail length (Supporting Information 2). We then averaged measurements across individuals within each OTU for downstream analyses. To test associations between migratory strategy and intraspecific morphological variation, we also calculated the coefficient of variation (mean/standard deviation) for each character and each taxon. This provides a scaled measure of variance for each character and allows testing whether migratory strategies differ in the amount of intraspecific variation.

We tested whether accounting for age (juvenile versus adult) and sex (female versus male) classes improved linear models explaining morphological measurements among taxa. We did this using the mulTree function of the mulTree package (Guillerme & Healy, 2020) in R, which assesses intraspecific variation while accounting for phylogenetic relatedness (e.g., Nations *et al.*, 2019). We found that including age class improved model fit for all characters except tarsus length and including sex class improved model fit for all characters except bill length and tarsus length (Supporting Information 1). We subsequently omitted juveniles from our analyses, but presented the results of both sexes combined in the main text because our results did not differ when analyzing sexes independently (Supporting Information 3).

### Phylogenetic Principal Components Analysis

We conducted a phylogenetic principal component analysis (PPCA) for bill measurements using the phyl.pca function from the phytools package (Revell, 2009) because this gives information on bill volume and shape that is not reflected in individual bill measurements (Table 1). Bill PPCA scores were included in the following phylogenetic analysis of variance (see next section). PPCA was also conducted for sexes separately (Supporting Information 3).

**Table 1.**
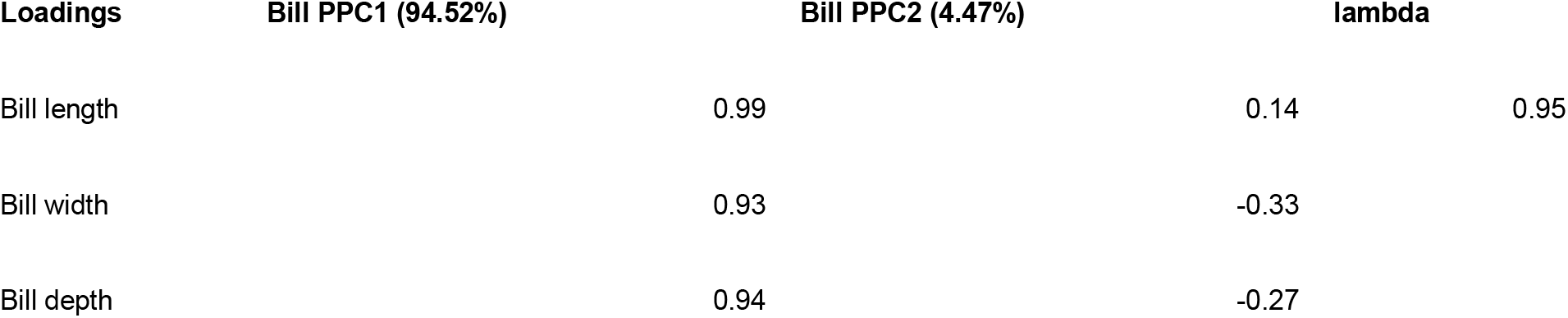
Results of phylogenetic principal component analysis (PPCA) of bill measurements. Percent variance explained by each eigenvector is in brackets for each principal component.

| Loadings | Bill PPC1 (94.52%) | Bill PPC2 (4.47%) | lambda |
| --- | --- | --- | --- |
| Bill length | 0.99 | 0.14 | 0.95 |
| Bill width | 0.93 | -0.33 |  |
| Bill depth | 0.94 | -0.27 |  |

### Phylogenetic Analysis of Variance

We compared morphology between migratory, partially migratory, and sedentary taxa via a phylogenetic analysis of variance (Garland *et al.*, 1993) with the phylANOVA function in the phytools package (Revell, 2012). As differences in body size can account for much of the variation among species (Albrecht, Gelvin, & Hartman, 1993; McCoy *et al.*, 2006; Revell, 2009; Berner, 2011), we extracted phylogenetic residuals (Revell, 2009) for each dependent variable (bill length, bill width, bill depth, wing chord, Kipp’s distance, and tail length) with tarsus length as an approximation of body size and the independent variable. We opted to use tarsus instead of mass to adjust for body size (Rising & Somers, 1989; Senar & Pascual, 1997) because mass can change seasonally (particularly in migratory birds; Lindström & Piersma, 1993) and also varies by sex in some kingbirds (e.g., *T. melancholicus*, Jahn *et al.*, 2010; *T. savana*, Carvalho Provinciato *et al.*, 2018; *T. tyrannus*, Murphy, 2007). To additionally support this decision, we used the mulTree function of the mulTree package to test whether tarsus length had the highest correlation among measurements while accounting for phylogeny, finding that tarsus length the highest correlation with body mass among all variables for the individuals that had mass data (Supporting Information 1). We then conducted a phylogenetic ANOVA for each character using the residual values (with 1000 replicates) and reported mean P-values for pairwise, post-hoc comparisons of mean values and coefficients of variation between migration categories. As such, we conducted 51 tests of statistical significance, which raises the issue of multiple hypothesis testing (Shaffer, 1995). Opinions differ on whether or not to adjust P-values when testing multiple hypotheses (Curran-Everett, 2000) and the 0.05 conventional threshold is ultimately arbitrary (Wasserstein, Schirm, & Lazar, 2019). Nonetheless, we report both uncorrected P-values and Bonferroni-corrected P-values (n = 3 tests for each set of pairwise comparisons of means or coefficients of variation). We also performed ancestral state reconstructions of wing chord length with the contMap function in the phytools package (Revell, 2012) and of migratory status with the fit.mle function in the diversitree package (FitzJohn, 2012). All statistical analyses were performed using program R version 4.0.3 (R Core Team, 2020).

## Results

The mean values for each morphometric character +/− standard deviation for all adults in each OTU are reported in Table S1.1. We found that migratory taxa had longer wing chords (Figure 2F, Table 2) and pointier wings than sedentary taxa (Figure 2D, Table 2). However, we found no association between migratory status and coefficients of variation in bill morphology (Figure 2G-I, Table 2). In agreement with our prediction, we found no relationship between migratory status and bill length, width, depth, or bill PPC1 (Figure 2A-C, Figure 3A, Table 2). However, migratory taxa had shorter, wider, and deeper bills (bill PPC2) compared to sedentary taxa, which had longer, shallower, and more narrow bills (Figure 3B, Table 2).

**Figure 2.**
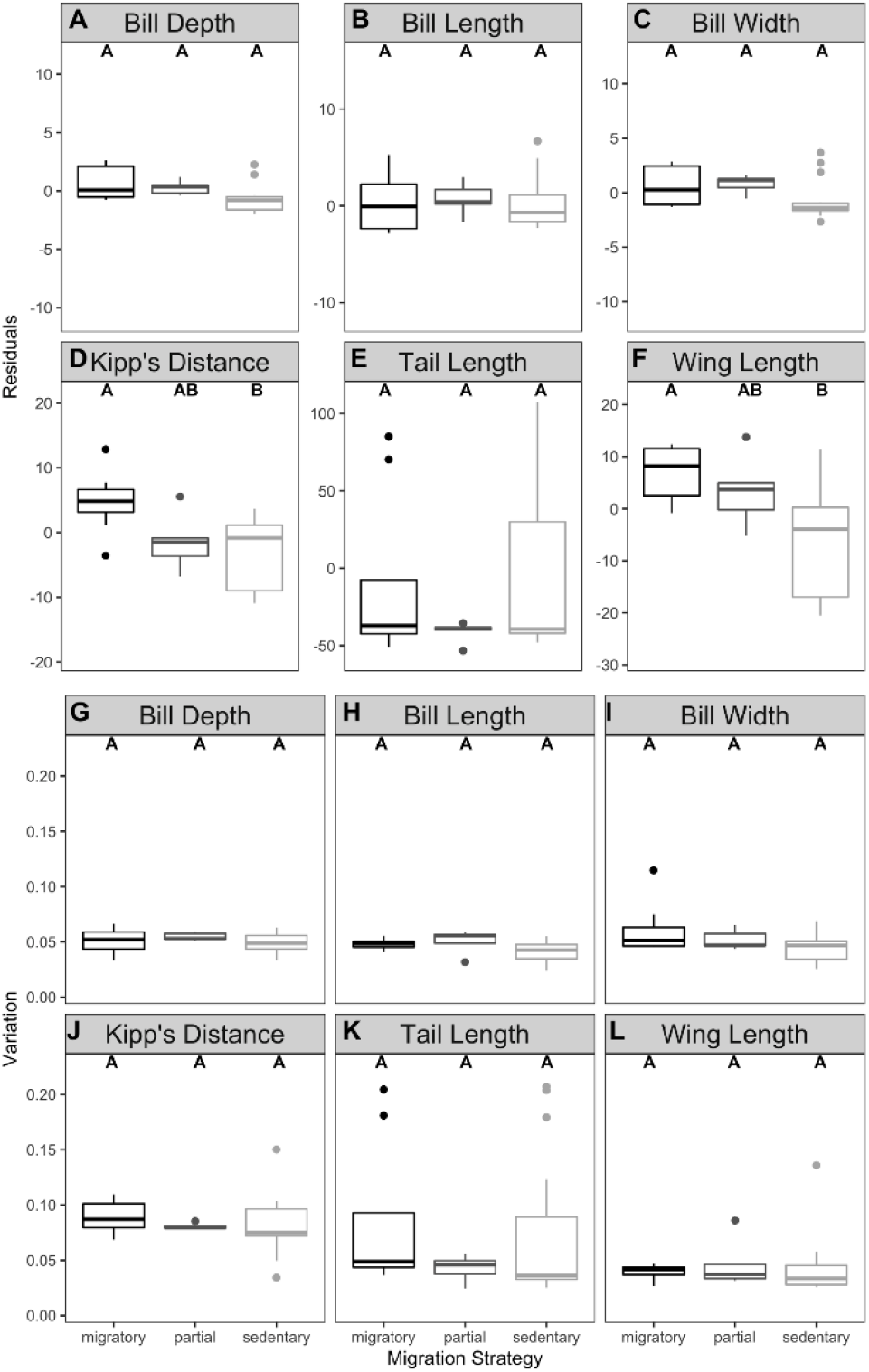
Phylogenetic ANOVA results comparing residuals of bill and feather morphometrics (A-F), and coefficient of variation in bill and feather morphometrics (G-L) across migratory, partially migratory, and sedentary kingbird OTUs. Note: although Kipp’s distance was significantly different between migratory and sedentary taxa for all adults, this was not significantly different after Bonferroni correction when males were assessed separately (Supporting Information Table 3.6). Letters above each barplot show groupings resulting from post-hoc pairwise comparisons.

**Table 2.**
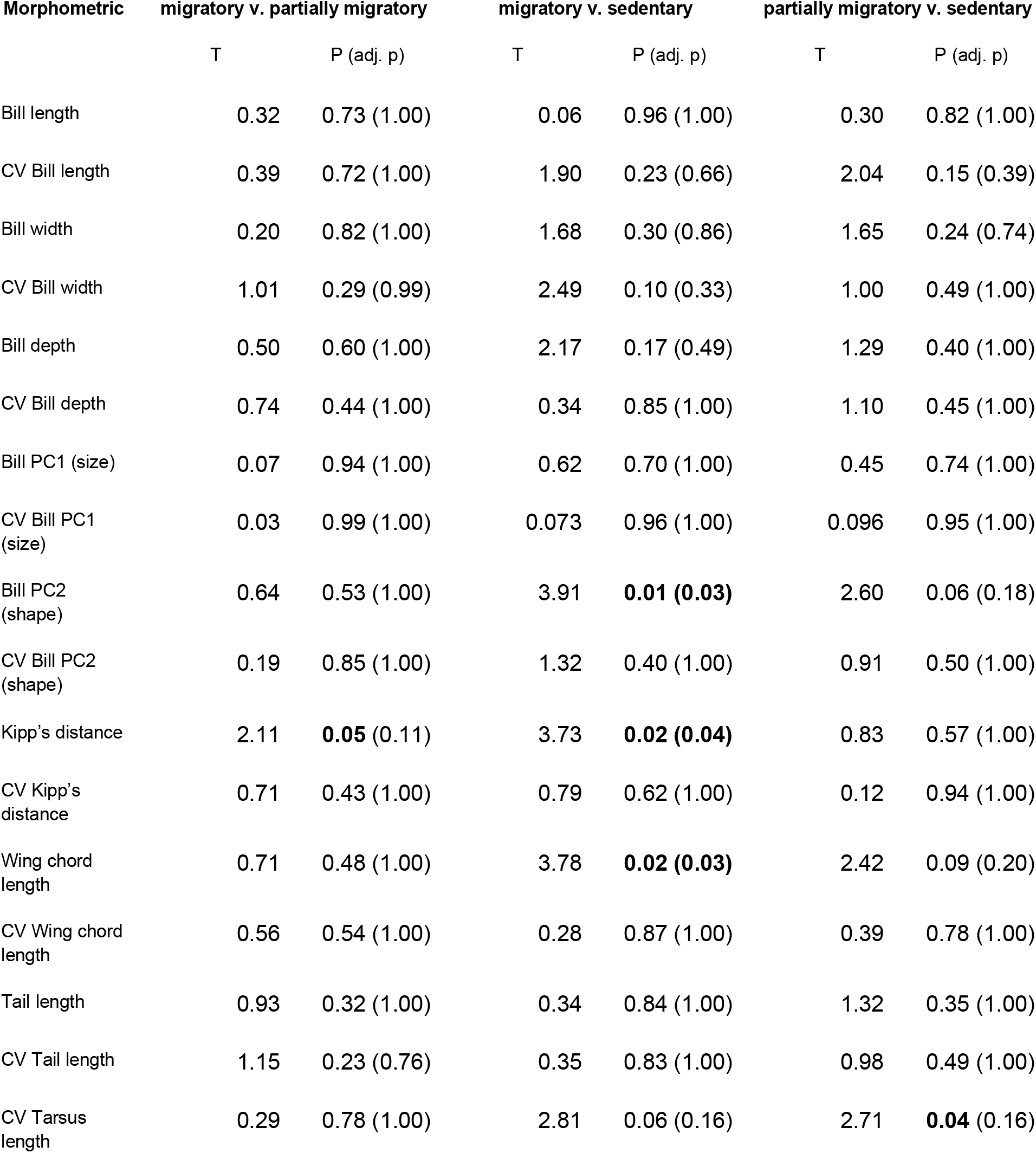
T and P values from phylogenetic ANOVA analysis of adult *Tyrannus* flycatchers. Values shown in brackets are P values with Bonferroni adjustment (n = 3) to account for multiple hypothesis testing. Significant results are in bold. Model F and Pr(>F) values can be found in Table S1.4.

**Figure 3.**
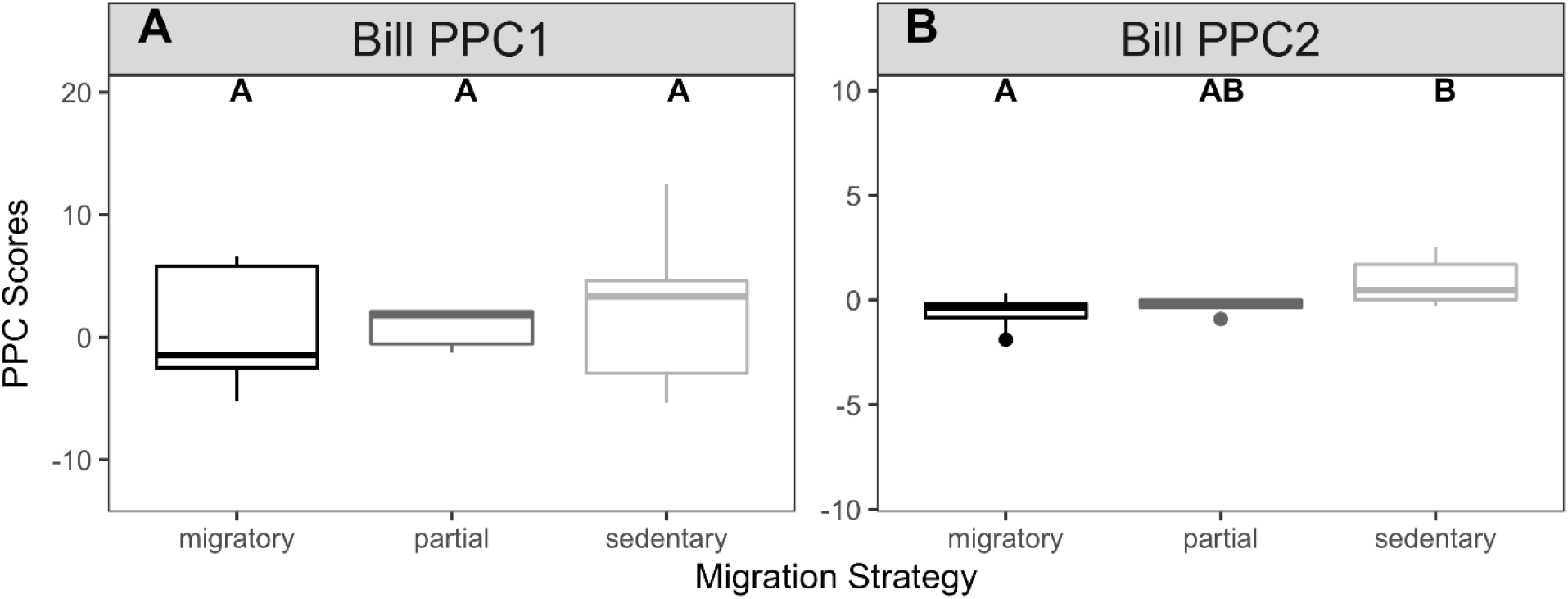
Phylogenetic ANOVA results comparing bill phylogenetic principal component (PPC) scores across For Peer Review migratory, partially migratory, and sedentary kingbird OTUs. Letters above each barplot show groupings resulting from post-hoc pairwise comparisons

## Discussion

We found evidence of an association between migratory status and wing morphology in the kingbirds. This corroborates the idea that migratory birds have evolved morphological features that help in sustained, long-distance flight, such as longer wings (Winkler & Leisler, 1992; Milá, Wayne, & Smith, 2008; Neto *et al.*, 2013; Carvalho Provinciato *et al.*, 2018). There is a strong genetic component determining wing length (Böhning-Gaese & Oberrath, 1999; Tarka *et al.*,2010), and among migratory taxa, wing length is positively associated with migratory distance (Förschler & Bairlein, 2011; Rönn, Shafer, & Wolf, 2016; Vágási *et al.*, 2016; Carvalho Provinciato *et al.*, 2018). Partial migration, wherein only some individuals migrate, is thought to be an evolutionary stepping-stone between sedentarism and obligate migration (Berthold, 1999; Bell, 2000; Chapman *et al.*, 2011). However, partial migration can also be sex and age-class dependent, as is the case for *T. melancholicus melancholicus* (Jahn *et al.*, 2010). Longer wings may allow for more flexibility in altering migration speeds according to conditions experienced during migration (Hahn *et al.*, 2016), but there may be ecological limits and trade-offs imposed on wing length by other selective pressures, like shorter wings for aerial agility among arboreal species (Sheard *et al.*, 2020), for evading predators (Fernández & Lank, 2007), or for hovering flight (Marchetti, Price, & Richman, 1995).

Our finding of differences in bill shape between migratory and sedentary taxa suggests that bill shape is a potential target of selection pressures related to migration ecology. However, our comparisons of coefficients of variation did not support intraspecific competition as a driver of diversifying selection in bill morphology among generalist migratory kingbirds. The ratio of bill length/bill width holds potential significance in foraging behaviors in other tyrannids (Botero-Delgadillo & Bayly, 2012), but our findings disagree with other research across the Tyrannidae describing generalists as having intermediate bill morphologies (Fitzpatrick, 1985). We found that sedentary taxa had longer, narrower, and shallower bills compared to migratory taxa (Figure 1B; Table 1). Previous work has shown that directional selection increasing bill length in dietary specialists may be driven by interspecific competition, including from closely related taxa (Freed, Medeiros, & Cann, 2016). Thus, other selective pressures may be acting upon the bill shape and size. For example, bill morphology has been linked with thermoregulation in dietary generalists, and bill width and depth are adaptive for dissipating heat during migration for improved thermoregulation (Danner & Greenberg, 2015; Friedman *et al.*, 2017; Danner *et al.*,2017). If convergence on bill shape is driven mainly by diet, future research integrating multivariate or nonlinear bill morphometrics (e.g., hooked bills, lateral and longitudinal bill curvature, bill surface area; Shao *et al.*, 2016) with degrees of intraguild dietary specialization and foraging behaviors would broaden our understanding of selection mechanisms shaping bill phenotypes.

The range of phenotypes shaped by migratory status is likely to be more varied than has been historically detectable using linear measurements of morphology alone. Divergence in bill shape between migratory and sedentary taxa may define an important component of the suite of co-adaptations for migratory life histories (Bell, 2000). Morphology may be driven by selection related to migration itself, such as an increased need for heat dissipation or conservation in flight(see above). Alternatively, morphology may be shaped by a more complex competitive landscape that affects migratory and sedentary taxa differently. In taxa occupying tropical regions, like many kingbirds, the dry season is when both intra- and interspecific competition increase due to ‘winter food limitation’ and competition with migrants returning to wintering grounds, and competition with juvenile conspecifics (Hespenheide, 1975; Emlen, 1977; Orejuela, Raitt, & Álvarez, 1980; Stiles, 1980; Waide, 1980; Rappole, Ramos, & Winker, 1989; Rappole, 1995; Sherry, 2005; Danner *et al.*, 2013). Selection may favor longer bills among sedentary taxa to resolve both intra- and interspecific conflict (Table 1, Figure 3B), allowing access to more temporally stable food resources in microhabitats buffered from spatial and temporal environmental fluctuations (Bell, 2011).

Migratory and sedentary taxa may not differ in linear, individual bill metrics if a combination of characteristics (e.g., Table 1, Figure 3B) support a more generalist foraging strategy. For example, specific bill shapes could accommodate spatiotemporal changes in the competitive landscape (Navalón *et al.*, 2019) or provide increased stability for prey capture in perch-gleaning or hover-gleaning foraging techniques (Fitzpatrick, 1985; Fitzpatrick *et al.*, 2004; Botero-Delgadillo, 2011; Botero-Delgadillo & Bayly, 2012). The diversity of habitats that migratory taxa encounter during their annual cycle may shape phenotypes in more complex ways than previously thought, and techniques that incorporate more advanced characterizations of morphological variation may expose novel insight into how movement life history strategies shape phenotypes (e.g., Pol *et al.*, 2009; Mallarino *et al.*, 2011; Navalón *et al.*, 2019; Alonso *et al.*, 2020; Medina *et al.*, 2020).

Aspects of both wing and bill morphology appear to have evolved in association with migratory status in kingbirds that have migratory, partially migratory, and sedentary taxa. Our results suggest that migratory status has shaped wing morphology in a widespread avian genus, and that multivariate bill shape metrics may differ between sedentary and migratory lineages. Thus, adaptive phenotypes may be related to migratory status in more complex ways than previously understood. Assessments of the various mechanisms driving patterns in bill shape (e.g., heat dissipation, foraging strategy, and competitive landscape) across a broader range of taxonomic groups that differ in migration strategies would complement our refinement of the morphology of migration in kingbirds.

## Data Availability

The data underlying the work, and the R code for all statistical analyses and results shared in tables and figures is available from the following public Github repository (https://github.com/mmacphe/Tyrannus_morphology). The voucher table and measurements of all individuals will also be made available at Dryad upon publication.

## Acknowledgements

This manuscript has been Accepted at the Biological Journal of the Linnean Society (August 02, 2021). We thank José Ignacio (“Nacho”) Giraldo Arango for measuring specimens, and Brenna and Michael Wells, and Shelbey Hearn for their assistance measuring specimens used in this study. Ben Marks, Shannon Hacket, and John Bates from the Field Museum of Natural History, Bradley Millen, and Allan Baker from the Royal Ontario Museum, Gary Stiles from the Instituto de Ciencias Naturales, Universidad Nacional de Colombia, Jacob Saucier, Helen James, and Gary Graves from the Smithsonian National Museum of Natural History, and Steven Cardiff, and Van Remsen from the Louisiana State University Museum of Natural Science provided assistance in accessing the specimens used in this study. Thanks to Robb Brumfield and Michael Harvey for providing an early-access version of the suboscine phylogeny for our study. We also thank Calandra Stanley, Caleb McMahan, the Toronto MacPhersons and the Bogotá Acevedos for their special help aiding the lead author to visit collections. Travel to measure specimens was supported in part from a scholar award from the James S. McDonnell Foundation. Samantha Rutledge, David Vander Pluym, and Subir Shakya provided helpful feedback on an earlier version of this manuscript, and we thank an anonymous reviewer and Prof John A. Allen for their insightful peer review. We wish to acknowledge www.gbif.org for providing the ROM database (accessed September 30, 2020) and the Field Museum database (accessed October 20, 2020) online that was used to validate locality data previously recorded by hand. Amethys E’etessam, Clare Lister, and Trey Hendrix also helped with georeferencing from locality data that was a part of this project.

## Supporting Information

Supporting Information 1 – Supplementary Results

Supporting Information 2 – ImageJ protocol for measuring long *Tyrannus* tail lengths

Supporting Information 3 – Sex-specific results: means for each morphometric, phylogenetic principal components analysis (PPCA), and phylogenetic ANOVA

## Literature Cited

Albrecht GH, Gelvin BR & Hartman SE. 1993. Ratios as a size adjustment in morphometrics. American Journal of Physical Anthropology 91: 441–468.

Alonso D, Fernández-Eslava B, Edelaar P & Arizaga J. 2020. Morphological divergence among Spanish Common Crossbill populations and adaptations to different pine species. Ibis 162: 1279–1291.

Bartoń K. 2020. MuMIn: multi-model inference.

Bell C. 2000. Process in the evolution of bird migration and pattern in avian ecogeography. Journal of Avian Biology 31: 258–265.

Bell C. 2011. Resource buffering and the evolution of bird migration. Evolutionary Ecology 25:91–106.

Berner D. 2011. Size correction in biology: how reliable are approaches based on (common) principal component analysis? Oecologia 166: 961–971.

Berthold P. 1999. A comprehensive theory for the evolution, control and adaptability of avian migration. Ostrich 70: 1–11.

Böhning-Gaese K & Oberrath R. 1999. Phylogenetic effects on morphological, life-history, behavioural and ecological traits of birds. : 18.

Bolnick DI, Svanbäck R, Araújo MS & Persson L. 2007. Comparative support for the niche variation hypothesis that more generalized populations also are more heterogeneous. Proceedings of the National Academy of Sciences 104: 10075–10079.

Botero-Delgadillo E. 2011. Cuantificando el comportamiento: Estrategias de búsqueda y ecología de forrajeo de 12 especies sintópicas de Atrapamoscas (Tyrannidae) en la Sierra Nevada de Santa Marta, Colombia. Revista Brasileira de Ornitologia 19: 343–357.

Botero-Delgadillo E & Bayly NJ. 2012. Does morphology predict behavior? Correspondence between behavioral and morphometric data in a Tyrant-flycatcher (Tyrannidae) assemblage in the Santa Marta Mountains, Colombia. Journal of Field Ornithology 83: 329–342.

Boyle WA. 2008. Partial migration in birds: tests of three hypotheses in a tropical lekking frugivore. Journal of Animal Ecology 77: 1122–1128.

Boyle WA & Conway CJ. 2007. Why Migrate? A Test of the Evolutionary Precursor Hypothesis. The American Naturalist 169: 344–359.

Carvalho Provinciato IC, Araújo MS & Jahn AE. 2018. Drivers of wing shape in a widespread Neotropical bird: a dual role of sex-specific and migration-related functions. Evolutionary Ecology 32: 379–393.

Chapman BB, Brönmark C, Nilsson JÅ & Hansson LA. 2011. The ecology and evolution of partial migration. Oikos 120: 1764–1775.

Chapman BB, Hulthén K, Brönmark C, Nilsson PA, Skov C, Hansson LA & Brodersen J. 2015. Shape up or ship out: migratory behaviour predicts morphology across spatial scale in a freshwater fish. Journal of Animal Ecology 84: 1187–1193.

Chesser RT & Levey DJ. 1998. Austral Migrants and the Evolution of Migration in New World Birds: Diet, Habitat, and Migration Revisited. The American Naturalist 152: 311–319.

Cintra R. 1997. Spatial Distribution and Foraging Tactics of Tyrant Flycatchers in Two Habitats in the Brazilian Amazon. Studies on Neotropical Fauna and Environment 32: 17–27.

Clegg SM & Owens PF. 2002. The ‘island rule’ in birds: medium body size and its ecological explanation. Proceedings of the Royal Society of London. Series B: Biological Sciences 269:1359–1365.

Clements JF, Schulenberg TS, Iliff MJ, Billerman SM, Fredericks TA, Sullivan BL & Wood CL. 2019. The eBird/Clements Checklist of Birds of the World: v2019.

Cooney CR, Bright JA, Capp EJR, Chira AM, Hughes EC, Moody CJA, Nouri LO, Varley ZK & Thomas GH. 2017. Mega-evolutionary dynamics of the adaptive radiation of birds. Nature 542: 344–347.

Cox GW. 1968. The Role of Competition in the Evolution of Migration. Evolution 22: 180–192.

Cox GW. 1985. The Evolution of Avian Migration Systems between Temperate and Tropical Regions of the New World. The American Naturalist 126: 451–474.

Crossin GT, Hinch SG, Farrell AP, Higgs DA, Lotto AG, Oakes JD & Healey MC. 2004. Energetics and morphology of sockeye salmon: effects of upriver migratory distance and elevation. Journal of Fish Biology 65: 788–810.

Danner RM, Greenberg RS, Danner JE, Kirkpatrick LT & Walters JR. 2013. Experimental support for food limitation of a short-distance migratory bird wintering in the temperate zone. Ecology 94: 2803–2816.

Danner RM, Gulson-Castillo ER, James HF, Dzielski SA, Frank DC III, Sibbald ET & Winkler DW. 2017. Habitat-specific divergence of air conditioning structures in bird bills. The Auk 134: 65–75.

Danner RM & Greenberg R. 2015. A critical season approach to Allen’s rule: Bill size declines with winter temperature in a cold temperate environment. Journal of Biogeography 42: 114–120.

Emlen JT. 1977. Land Bird Communities of Grand Bahama Island: The Structure and Dynamics of an Avifauna. Ornithological Monographs: iii–129.

Felice RN, Tobias JA, Pigot AL & Goswami A. 2019. Dietary niche and the evolution of cranial morphology in birds. Proceedings of the Royal Society B: Biological Sciences 286: 20182677.

Fernández G & Lank DB. 2007. Variation in the Wing Morphology of Western Sandpipers (Calidris Mauri) in Relation to Sex, Age Class, and Annual Cycle. The Auk 124: 1037–1046.

Fiedler W. 2005. Ecomorphology of the External Flight Apparatus of Blackcaps ( Sylvia atricapilla ) with Different Migration Behavior. Annals of the New York Academy of Sciences 1046: 253–63.

FitzJohn RG. 2012. Diversitree: comparative phylogenetic analyses of diversification in R. Methods in Ecology and Evolution 3: 1084–1092.

Fitzpatrick JW. 1981. Search strategies of tyrant flycatchers. Animal Behaviour 29: 810–821.

Fitzpatrick JW. 1985. Form, foraging behavior, and adaptive radiation in the Tyrannidae BT - Neo-Tropical Ornithology Ornithological Monographs No. 26. In: Neo-Tropical Ornithology Ornithological Monographs No. 26., 447–470.

Fitzpatrick JW, Bates JM, Bostwick KS, Caballero IC, Clock BM, Farnsworth A, Hosner PA, Joseph L, Langham GM & Lebbin DJ. 2004. Family Tyrannidae (tyrant-flycatchers). Handbook of the birds of the world 9: 170–462.

Fitzpatrick W & Schauensee D. 1980. Foraging behavior of Tyrant flycatchers. : 43–57.

Förschler MI & Bairlein F. 2011. Morphological Shifts of the External Flight Apparatus across the Range of a Passerine (Northern Wheatear) with Diverging Migratory Behaviour. PLOS ONE 6: e18732.

Freed LA, Medeiros MCI & Cann RL. 2016. Multiple Reversals of Bill Length over 1.7 Million Years in a Hawaiian Bird Lineage. The American Naturalist 187: 363–371.

Friedman NR, Harmáčková L, Economo EP & Remeš V. 2017. Smaller beaks for colder winters: Thermoregulation drives beak size evolution in Australasian songbirds. Evolution 71:2120–2129.

Gabriel V de A & Pizo MA. 2005. Foraging behavior of tyrant flycatchers (Aves, Tyrannidae) in Brazil. Revista Brasileira de Zoologia 22: 1072–1077.

Garland T, Dickerman AW, Janis CM & Jones JA. 1993. phylogenetic analysis of covariance by computer simulation. Systematic Biology 42: 265–292.

Gómez C, Tenorio EA, Montoya P & Cadena CD. 2016. Niche-tracking migrants and niche-switching residents: evolution of climatic niches in New World warblers (Parulidae). Proceedings of the Royal Society B: Biological Sciences 283: 20152458.

Gómez-Bahamón V, Márquez R, Jahn AE, Miyaki CY, Tuero DT, Laverde-R O, Restrepo S & Cadena CD. 2020a. Speciation Associated with Shifts in Migratory Behavior in an Avian Radiation. Current Biology 30: 1312–1321.e6.

Gómez-Bahamón V, Tuero DT, Castaño MI, Jahn AE, Bates JM & Clark CJ. 2020b. Sonations in Migratory and Non-migratory Fork-tailed Flycatchers ( *Tyrannus savana* ). Integrative and Comparative Biology 60: 1147–1159.

Guillerme T & Healy K. 2020. mulTree: performs MCMCglmm on Multiple Phylogenetic Trees.

Haberman K, MacKenzie DI & Rising JD. 1991. Geographic Variation in the Gray Kingbird (Variación geográfica en Tyrannus dominicensis). Journal of Field Ornithology 62: 117–131.

Hahn S, Korner-Nievergelt F, Emmenegger T, Amrhein V, Csörgő T, Gursoy A, Ilieva M, Kverek P, Pérez-Tris J, Pirrello S, Zehtindjiev P & Salewski V. 2016. Longer wings for faster springs – wing length relates to spring phenology in a long-distance migrant across its range. Ecology and Evolution 6: 68–77.

Harvey M, Bravo G, Claramunt S, Cuervo A, Derryberry G, Battilana J, Seeholzer G, McKay JS, O’Meara BC, Faircloth BC, Edwards S, Pérez-Emán J, Moyle RG, Sheldon FH, Aleixo A, Smith BT, Chesser RT, Silveira LF, Cracraft J, Brumfield RT & Derryberry EP. 2020. The evolution of a tropical biodiversity hotspot. Science 370: 1343–1348.

Herrera CM. 1978. Ecological Correlates of Residence and Non-Residence in a Mediterranean Passerine Bird Community. Journal of Animal Ecology 47: 871–890.

Hespenheide HA. 1975. Selective Predation by Two Swifts and a Swallow in Central America. Ibis 117: 82–99.

Hromada M & Tryjanowski P. 2003. Animals of different phenotype differentially utilise dietary niche - the case of the Great Grey Shrike Lanius excubitor. 80: 8.

Jahn AE, Levey DJ, Mamani AM, Saldias M, Alcoba A, Ledezma MJ, Flores B, Vidoz JQ & Hilarion F. 2010a. Seasonal differences in rainfall, food availability, and the foraging behavior of Tropical Kingbirds in the southern Amazon Basin. Journal of Field Ornithology 81: 340–348.

Jahn AE, Levey DJ, Hostetler JA & Mamani AM. 2010b. Determinants of partial bird migration in the Amazon Basin. Journal of Animal Ecology 79: 983–992.

Jahn AE & Tuero DT. 2020. Fork-tailed Flycatcher (Tyrannus savana). Birds of the World.

Johansson F, Söderquist M & Bokma F. 2009. Insect wing shape evolution: independent effects of migratory and mate guarding flight on dragonfly wings. Biological Journal of the Linnean Society 97: 362–372.

Kipp F. 1942. Über Flügelbau und Wanderzug der Vögel. Biologisches Zentralblatt 62: 289–299.

Kipp FA. 1958. Zur geschichte des Vogelzuges auf der grundlage der Flügelanpassungen. Vogelwarte 19: 233–242.

Knudsen R, Siwertsson A, Adams CE, Garduño-Paz M, Newton J & Amundsen PA. 2011. Temporal stability of niche use exposes sympatric Arctic charr to alternative selection pressures. Evolutionary Ecology 25: 589–604.

Lack D. 1971. Ecological isolation in birds.

Leisler B. 1990. Selection and Use of Habitat of Wintering Migrants. In: Gwinner E, ed. Bird Migration. Berlin, Heidelberg: Springer Berlin Heidelberg, 156–174.

Levey DJ & Stiles FG. 1992. Evolutionary Precursors of Long-Distance Migration: Resource Availability and Movement Patterns in Neotropical Landbirds. The American Naturalist 140:447–476.

Lindström Å & Piersma T. 1993. Mass changes in migrating birds: the evidence for fat and protein storage re-examined. Ibis 135: 70–78.

Lockwood R, Swaddle JP & Rayner JMV. 1998. Avian Wingtip Shape Reconsidered: Wingtip Shape Indices and Morphological Adaptations to Migration. Journal of Avian Biology 29: 273–292.

Mallarino R, Grant PR, Grant BR, Herrel A, Kuo WP & Abzhanov A. 2011. Two developmental modules establish 3D beak-shape variation in Darwin’s finches. Proceedings of the National Academy of Sciences 108: 4057–4062.

Marchetti K, Price T & Richman A. 1995. Correlates of Wing Morphology with Foraging Behaviour and Migration Distance in the Correlates of wing morphology with foraging behaviour and migration distance in the genus Phylloscopus. Journal of Avian Biology 26: 177–181.

McCoy MW, Bolker BM, Osenberg CW, Miner BG & Vonesh JR. 2006. Size correction: comparing morphological traits among populations and environments. Oecologia 148: 547–554.

Medina JJ, Maley JM, Sannapareddy S, Medina NN, Gilman CM & McCormack JE. 2020. A rapid and cost-effective pipeline for digitization of museum specimens with 3D photogrammetry. PLOS ONE 15: e0236417.

Milá B, Wayne RK & Smith TB. 2008. Ecomorphology of Migratory and Sedentary Populations of the Yellow-Rumped Warbler (Dendroica Coronata). The Condor 110: 335–344.

Minias P, Meissner W, Włodarczyk R, Ożarowska A, Piasecka A, Kaczmarek K & Janiszewski T. 2015. Wing shape and migration in shorebirds: a comparative study. Ibis 157: 528–535.

Mobley JA. 2002. Molecular phylogenetics and the evolution of nest building in kingbirds and their allies (Aves: Tyrannidae). University of California, Berkeley.

Mobley JA & de Juana E. 2020. Loggerhead Kingbird (Tyrannus caudifasciatus). Birds of the World.

Mönkkönen M. 1995. Do migrant birds have more pointed wings?: A comparative study. Evolutionary Ecology 9: 520–528.

Moreau RE. 1952. The Place of Africa in the Palaearctic Migration System. Journal of Animal Ecology 21: 250–271.

Morse DH. 1971. The Insectivorous Bird as an Adaptive Strategy. Annual Review of Ecology and Systematics 2: 177–200.

Mulvihill RS & Chandler CR. 1990. The Relationship between Wing Shape and Differential Migration in the Dark-Eyed Junco. The Auk 107: 490–499.

Murphy MT. 2007. A Cautionary Tale: Cryptic Sexual Size Dimorphism in a Socially Monogamous Passerine. The Auk 124: 515–525.

Nations JA, Heaney LR, Demos TC, Achmadi AS, Rowe KC & Esselstyn JA. 2019. A simple skeletal measurement effectively predicts climbing behaviour in a diverse clade of small mammals. Biological Journal of the Linnean Society 128: 323–336.

Navalón G, Bright JA, Marugán-Lobón J & Rayfield EJ. 2019. The evolutionary relationship among beak shape, mechanical advantage, and feeding ecology in modern birds*. Evolution 73:422–435.

Neto JM, Gordinho L, Belda EJ, Marín M, Monrós JS, Fearon P & Crates R. 2013. Phenotypic Divergence among West European Populations of Reed Bunting Emberiza schoeniclus: The Effects of Migratory and Foraging Behaviours. PLOS ONE 8: e63248.

Norberg UM. 1990. Vertebrate flight: mechanics, physiology, morphology, ecology and evolution. Berlin: Springer.

Norberg UM. 1995. WING DESIGN AND MIGRATORY FLIGHT. Israel Journal of Ecology and Evolution 41: 297–305.

Orejuela JE, Raitt RJ & Álvarez H. 1980. Differential use by North American migrants of three types of Colombian forests: 253-264 (en) Keast, A. & Morton, ES (eds.). Migrant Birds in the Neotropics: Ecology, Behavior, Distribution and Conservation. Smithsonian Institution Press. Washington DC, USA.

Pennycuick C. 2008. The membrane wings of bats and pterosaurs Modelling the Flying Bird ed CJ Pennycuick.

Pérez-Tris J & Tellería JL. 2001. Age-related variation in wing shape of migratory and sedentary Blackcaps Sylvia atricapilla. Journal of Avian Biology 32: 207–213.

Pol MVD, Ens BJ, Oosterbeek K, Brouwer L, Verhulst S, Tinbergen JM, Rutten AL & Jong MD. 2009. Oystercatchers’ Bill Shapes as a Proxy for Diet Specialization: More Differentiation than Meets the Eye. Ardea 97: 335–347.

Pyle P, Howell SNG, Institute for Bird Populations, & Point Reyes Bird Observatory. 1997. Identification guide to North American birds: a compendium of information on identifying, ageing, and sexing ‘near-passerines’ and passerines in the hand. Part I, Part I,. Bolinas, Calif.: Slate Creek Press.

R Core Team. 2020. R: A Language and Environment for Statistical Computing (4.0.3)[Computer software]. R Foundation for Statistical Computing. Retrieved from http://www.R-proje ....

Rappole JH. 1995. The ecology of migrant birds: a neotropical perspective. Washington; London: Smithsonian institution Press.

Rappole JH, Ramos MA & Winker K. 1989. Wintering Wood Thrush Movements and Mortality in Southern Veracruz. The Auk 106: 402–410.

Rayner JMV. 1988. The evolution of vertebrate flight. Biological Journal of the Linnean Society 34: 269–287.

Reaney AM, Bouchenak-Khelladi Y, Tobias JA & Abzhanov A. 2020. Ecological and morphological determinants of evolutionary diversification in Darwin’s finches and their relatives. Ecology and Evolution 10: 14020–14032.

Regosin JV & Pruett-Jones S. 2001. Sexual Selection and Tail-Length Dimorphism in Scissor-Tailed Flycatchers. The Auk 118: 167–175.

Revell LJ. 2009. Size-Correction and Principal Components for Interspecific Comparative Studies. Evolution 63: 3258–3268.

Revell LJ. 2012. phytools: an R package for phylogenetic comparative biology (and other things). Methods in Ecology and Evolution 3: 217–223.

Rising JD & Somers KM. 1989. The Measurement of Overall Body Size in Birds. The Auk 106: 666–674.

Rönn JAC von, Shafer ABA & Wolf JBW. 2016. Disruptive selection without genome-wide evolution across a migratory divide. Molecular Ecology 25: 2529–2541.

Senar JC & Pascual J. 1997. Keel and tarsus length may provide a good predictor of avian body size. Ardea 85.

Shao S, Quan Q, Cai T, Song G, Qu Y & Lei F. 2016. Evolution of body morphology and beak shape revealed by a morphometric analysis of 14 Paridae species. Frontiers in Zoology 13: 30.

Sheard C, Neate-Clegg MHC, Alioravainen N, Jones SEI, Vincent C, MacGregor HEA, Bregman TP, Claramunt S & Tobias JA. 2020. Ecological drivers of global gradients in avian dispersal inferred from wing morphology. Nature Communications 11: 2463.

Sherry TW. 1984. Comparative Dietary Ecology of Sympatric, Insectivorous Neotropical Flycatchers (Tyrannidae). Ecological Monographs 54: 313–338.

Sherry TW. 2005. Does winter food limit populations of migratory birds? Birds of two worlds: the ecology and evolution of migration: 414–425.

Snow DW. 1953. Systematics and comparative ecology of the genus Parus in the Palaearctic region.

Snow DW. 1954. The habitats of Eurasian tits (Parus spp.). Ibis 96: 565–585.

Stiles FG. 1980. Evolutionary implications of habitat relations between permanent and winter resident landbirds in Costa Rica. Migrant Birds in the Neotropics (A. Keast and ES Morton, Eds.) Smithsonian Institution Press, Washington DC: 421–435.

Svanbäck R & Schluter D. 2012. Niche Specialization Influences Adaptive Phenotypic Plasticity in the Threespine Stickleback. The American Naturalist 180: 50–59.

Swanson MT, Oliveros CH & Esselstyn JA. 2019. A phylogenomic rodent tree reveals the repeated evolution of masseter architectures. Proceedings of the Royal Society B: Biological Sciences 286: 20190672.

Tarka M, Åkesson M, Beraldi D, Hernández-Sánchez J, Hasselquist D, Bensch S & Hansson B. 2010. A strong quantitative trait locus for wing length on chromosome 2 in a wild population of great reed warblers. Proceedings of the Royal Society B: Biological Sciences 277: 2361–2369.

Tellería LJ, Blázquez M, De La Hera I & Pérez-Tris J. 2013. Migratory and resident Blackcaps Sylvia atricapilla wintering in southern Spain show no resource partitioning. Ibis 155: 750–761.

Tellería JL & Carbonell R. 1999. Morphometric Variation of Five Iberian Blackcap Sylvia atricapilla Populations. Journal of Avian Biology 30: 63–71.

Thiollay JM. 1988. Comparative Foraging Success of Insectivorous Birds in Tropical and Temperate Forests: Ecological Implications. Oikos 53: 17–30.

Vágási CI, Pap PL, Vincze O, Osváth G, Erritzøe J & Møller AP. 2016. Morphological Adaptations to Migration in Birds. Evolutionary Biology 43: 48–59.

Van Dijk A, Nakamura G, Rodrigues AV, Maestri R & Duarte L. 2021. Imprints of tropical niche conservatism and historical dispersal in the radiation of Tyrannidae (Aves: Passeriformes). Biological Journal of the Linnean Society.

Van Valen L. 1965. Morphological Variation and Width of Ecological Niche. The Amercian Naturalist 99: 377–390.

Voelker G. 2001. Morphological correlates of migratory distance and flight display in the avian genus Anthus. Biological Journal of the Linnean Society 73: 425–435.

Waide RB. 1980. Resource partitioning between migrant and resident birds: the use of irregular resources. Migrant birds in the Neotropics: ecology, behavior, distribution, and conservation: 337–352.

Wang X & Clarke JA. 2015. The evolution of avian wing shape and previously unrecognized trends in covert feathering. Proceedings of the Royal Society B: Biological Sciences 282: 20151935.

Wiedenfeld DA. 1991. Geographical Morphology of Male Yellow Warblers. The Condor 93:712–723.

Willis EO. 1974. Populations and Local Extinctions of Birds on Barro Colorado Island, Panama. Ecological Monographs 44: 153–169.

Wilson EO. 1961. The Nature of the Taxon Cycle in the Melanesian Ant Fauna. The American Naturalist 95: 169–193.

Winkler H & Leisler B. 1992. On the ecomorphology of migrants. Ibis 134: 21–28.

Winquist T & Lemon RE. 1994. Sexual Selection and Exaggerated Male Tail Length in Birds. The American Naturalist 143: 95–116.

Závorka L, Larranaga N, Lovén Wallerius M, Näslund J, Koeck B, Wengström N, Cucherousset J & Johnsson JI. 2020. Within-stream phenotypic divergence in head shape of brown trout associated with invasive brook trout. Biological Journal of the Linnean Society 129: 347–355.

