## Supporting Information 1 for "Morphology of migration: Associations between wing, and bill morphology and migration in kingbirds (*Tyrannus*)"

**Supporting Information 1** - Supplementary Results

**Table S1.1**. Morphological measurement means for adults of 28 *Tyrannus* OTUs (millimeters). All measurements are shown ± standard deviation and with the sample size in brackets. After *Tyrannus savana monachus*, "CA" refers to Central America and "SA" refers to South America.

| OTU | Mass (g) | Bill length | Bill width | Bill depth | Tarsus length | Kipp's distance | Wing chord length | Tail length |
|---|---|---|---|---|---|---|---|---|
| *Tyrannus albogularis* | 38.29 ± 3.53 (17) | 15.14 ± 0.74 (36) | 8.61 ± 0.48 (35) | 6.37 ± 0.37 (33) | 17.58 ± 0.75 (36) | 28.52 ± 2.29 (26) | 105.09 ± 4.86 (26) | 91.02 ± 5.79 (26) |
| *Tyrannus caudifasciatus bahamensis* | NA | 20.19 ± 0.8 (13) | 9.45 ± 0.43 (14) | 7.52 ± 0.34 (14) | 22.3 ± 0.41 (14) | 25.34 ± 1.71 (12) | 107.99 ± 3.61 (12) | 84.94 ± 2.86 (12) |
| *Tyrannus caudifasciatus caudifasciatus* | NA | 19.23 ± 1.96 (77) | 8.82 ± 0.77 (76) | 7.21 ± 0.47 (62) | 22.46 ± 0.7 (78) | 23.35 ± 2.2 (61) | 103.44 ± 2.79 (62) | 85.16 ± 2.87 (53) |
| *Tyrannus caudifasciatus caymanensis* | 43.32 ± 5.57 (6) | 21.63 ± 0.93 (12) | 9.42 ± 0.25 (11) | 7.41 ± 0.29 (11) | 22.6 ± 0.62 (15) | 21.81 ± 1.84 (11) | 103.13 ± 2.77 (11) | 87.96 ± 3.77 (10) |
| *Tyrannus caudifasciatus flavescens* | NA | 19.95 ± 0.67 (21) | 8.47 ± 0.22 (21) | 7.18 ± 0.24 (21) | 22.61 ± 0.47 (21) | 23.51 ± 1.17 (21) | 104.74 ± 3.24 (21) | 83.9 ± 2.41 (21) |
| *Tyrannus caudifasciatus gabbii* | NA | 18.28 ± 0.64 (31) | 8.89 ± 0.4 (32) | 6.7 ± 0.36 (29) | 20.97 ± 0.72 (32) | 22.48 ± 1.63 (23) | 103.07 ± 3.59 (24) | 80.71 ± 2.58 (22) |
| *Tyrannus caudifasciatus jamaicensis* | 38.5 ± NA (1) | 20.22 ± 1.23 (16) | 9.11 ± 0.3 (15) | 7.07 ± 0.42 (14) | 21.79 ± 0.53 (16) | 22.98 ± 1.89 (12) | 100.65 ± 2.81 (12) | 83.59 ± 3.28 (12) |
| *Tyrannus caudifasciatus taylori* | 52.96 ± 0.75 (3) | 20.38 ± 0.85 (75) | 10.02 ± 0.5 (76) | 7.91 ± 0.32 (60) | 23.65 ± 0.7 (76) | 24.4 ± 1.67 (67) | 112.59 ± 3.29 (67) | 92.19 ± 2.46 (63) |
| *Tyrannus couchii* | 46.43 ± 6.98 (27) | 16.47 ± 1.77 (91) | 10.17 ± 0.84 (89) | 7.63 ± 0.46 (84) | 19.46 ± 0.76 (93) | 31.19 ± 2.5 (64) | 116.08 ± 5.07 (64) | 93.86 ± 5.13 (52) |
| *Tyrannus crassirostris crassirostris* | 56.36 ± 1.76 (5) | 19.93 ± 1.14 (22) | 12.5 ± 0.55 (22) | 10.36 ± 0.41 (21) | 20.36 ± 0.69 (24) | 36.46 ± 3.48 (20) | 127.05 ± 5.15 (20) | 95.37 ± 4.01 (15) |
| *Tyrannus crassirostris pompalis* | NA | 20.28 ± 0.89 (49) | 12.32 ± 0.72 (50) | 10.56 ± 0.49 (48) | 20.79 ± 0.77 (50) | 37.64 ± 3.33 (40) | 129.41 ± 3.58 (40) | 95.11 ± 3.48 (43) |
| *Tyrannus cubensis* | NA | 26.13 ± 0.72 (33) | 14.18 ± 0.48 (33) | 11.72 ± 0.48 (34) | 23.02 ± 0.76 (34) | 33.78 ± 2.57 (24) | 129.21 ± 3.71 (24) | 96.09 ± 5.19 (17) |

| | | | | | | | | |
|---|---|---|---|---|---|---|---|---|
| *Tyrannus dominicensis dominicensis* | 46.69 ± 7.88 (15) | 20.42 ± 1.42 (102) | 10.96 ± 0.63 (99) | 8.55 ± 0.64 (94) | 18.73 ± 0.61 (109) | 33.1 ± 2.54 (77) | 112.65 ± 3.7 (76) | 86.28 ± 3.4 (62) |
| *Tyrannus dominicensis vorax* | 50.56 ± 2.89 (9) | 22.82 ± 1.17 (63) | 12.63 ± 0.65 (61) | 9.33 ± 0.58 (56) | 19.34 ± 0.7 (69) | 31.29 ± 2.36 (50) | 114.3 ± 3.76 (50) | 89.4 ± 3.61 (34) |
| *Tyrannus forficatus* | 40.96 ± 5.3 (34) | 13.49 ± 0.62 (86) | 7.64 ± 0.36 (88) | 6.23 ± 0.32 (85) | 19.07 ± 0.75 (95) | 42.47 ± 5.92 (77) | 118.29 ± 6.27 (77) | 181.93 ± 47.89 (85) |
| *Tyrannus melancholicus despotes* | NA | 17.48 ± 0.91 (55) | 9.64 ± 0.56 (57) | 7.45 ± 0.56 (45) | 17.76 ± 0.98 (58) | 30.78 ± 2.86 (38) | 108.03 ± 9.07 (38) | 90.65 ± 4.58 (21) |
| *Tyrannus melancholicus melancholicus* | 41.04 ± 5.36 (72) | 17.37 ± 1.32 (253) | 9.7 ± 0.54 (251) | 7.45 Â± 0.6 (244) | 18.18 ± 0.78 (255) | 30.38 ± 2.68 (194) | 112.32 ± 4 (195) | 91.79 ± 4.81 (126) |
| *Tyrannus melancholicus satrapa* | 41.83 ± 5.2 (74) | 17.59 ± 1.11 (195) | 9.93 ± 0.58 (199) | 7.31 ± 0.42 (195) | 19.19 ± 0.9 (204) | 28.4 ± 2.31 (143) | 110.88 ± 4.36 (143) | 90.67 ± 4.73 (114) |
| *Tyrannus niveigularis* | 34.44 ± 3.03 (17) | 14.67 ± 0.54 (21) | 8.6 ± 0.5 (22) | 6.14 ± 0.34 (22) | 18.22 ± 0.79 (22) | 24.91 ± 2.03 (20) | 101.58 ± 3.58 (20) | 77.12 ± 2.37 (21) |
| *Tyrannus savana circumdatus* | NA | 12.15 ± 0.61 (25) | 6.37 ± 0.32 (25) | 5.19 ± 0.26 (25) | 17.62 ± 0.4 (25) | 33.77 ± 3.42 (23) | 104 ± 4.14 (23) | 221.53 ± 26.77 (20) |
| *Tyrannus savana monachus CA* | 27.6 ± 2.58 (11) | 11.23 ± 0.48 (57) | 6.49 ± 0.33 (55) | 5.23 ± 0.33 (54) | 17.43 ± 0.69 (58) | 32.18 ± 2.84 (32) | 100.98 ± 4.92 (32) | 232.56 ± 56.05 (20) |
| *Tyrannus savana monachus SA* | 26.97 ± 1.64 (6) | 11.55 ± 0.71 (97) | 6.35 ± 0.43 (96) | 5.34 ± 0.32 (85) | 17.35 ± 0.65 (45) | 32.93 ± 2.83 (36) | 100.6 ± 12.63 (76) | 215.54 ± 53.3 (59) |
| *Tyrannus savana sanctaemartae* | NA | 11.98 ± 0.37 (14) | 6.15 ± 0.24 (14) | 5.31 ± 0.24 (14) | 17.56 ± 0.57 (14) | 34.73 ± 3.36 (8) | 102.06 ± 5.69 (9) | 234.44 ± 47.77 (9) |
| *Tyrannus savana savana* | 28.82 ± 4.19 (22) | 11.11 ± 0.66 (138) | 6.5 ± 0.5 (139) | 5.28 ± 0.42 (123) | 17.72 ± 1.01 (110) | 35.21 ± 3.8 (66) | 104.97 ± 11.47 (93) | 202.94 ± 49.1 (78) |
| *Tyrannus tyrannus* | 40.89 ± 7.41 (50) | 13.47 ± 1.02 (108) | 8.05 ± 0.81 (106) | 6.31 ± 0.57 (105) | 18.83 ± 0.8 (108) | 37.25 ± 2.98 (72) | 114.44 ± 4.94 (73) | 78.73 ± 4.07 (62) |
| *Tyrannus verticalis* | 42.75 ± 3.76 (15) | 13.25 ± 0.95 (98) | 7.72 ± 0.46 (98) | 6.53 ± 0.45 (95) | 19.49 ± 0.76 (106) | 38.91 ± 4.08 (95) | 122.51 ± 5.75 (96) | 87.52 ± 4.43 (94) |
| *Tyrannus vociferans vociferans* | 46.18 ± 3.57 (44) | 15.01 ± 1.04 (182) | 8.63 ± 0.51 (183) | 6.96 ± 0.5 (173) | 19.79 ± 0.73 (198) | 37.27 ± 3.34 (117) | 126.74 ± 10.51 (118) | 89.56 ± 3.41 (105) |

| | | | | | | | | |
|---|---|---|---|---|---|---|---|---|
| *Tyrannus vociferans xenopterus* | NA | 14.74 ± 0.73 (12) | 8.36 ± 0.42 (12) | 6.62 ± 0.36 (12) | 19.8 ± 0.61 (12) | 35.99 ± 5.41 (6) | 125.09 ± 7.21 (6) | 90.63 ± 3.27 (7) |

**Table S1.2.** Tests of whether age or sex classes play a role in morphological measurements. We used the mulTree function in the mulTree library to test the role of age and sex classes on each morphological measurement.

| Morphological Measurement | Effect of Age Class (p-value) | Effect of Sex Class (p-value) |
|---|---|---|
| Bill Length | <0.01 | 0.07 |
| Bill Width | <0.01 | <0.01 |
| Bill Depth | <0.01 | <0.01 |
| Kipp's distance | <0.01 | <0.01 |
| Wing Chord Length | <0.01 | <0.01 |
| Tail Length | <0.01 | <0.01 |
| Tarsus Length | 0.24 | 0.36 |

**Table S1.3.** Test of whether tarsus length is the best morphological measurement to approximate body mass. We used the mulTree function in the mulTree library to assess the correlation of all measured morphologies with body mass for 438 individuals that had mass data on specimen tags (representing 18 taxa) while accounting for phylogeny.

| Coefficients | P value |
|---|---|
| (Intercept) | 0.0061 |
| Tarsus length | 0.00121 |
| Wing chord length | 0.00162 |
| Bill depth | 0.02747 |
| Kipp's distance | 0.04101 |
| Bill width | 0.52747 |
| Bill length | 0.56747 |
| Tail length | 0.81778 |

**Table S1.4**. F and Pr(>F) values from phylogenetic ANOVA. Adjusted P-value after Bonferroni correction for multiple hypothesis testing is only shown for the instance using polytomies added with the most recent common ancestor at 0.5 million years ago.

| Character of Interest | Most recent common ancestor 0.5 mya | | Most recent common ancestor 0.2 mya | | Most recent common ancestor 0.97 mya | |
|---|---|---|---|---|---|---|
| | F value | Pr(>F) value (adj. p) | F value | Pr(>F) value | F value | Pr(>F) value |
| Bill Length | 0.06 | 0.96 (0.96) | 0.06 | 0.96 | 0.06 | 0.95 |
| Bill Width | 2.14 | 0.32 (0.32) | 1.87 | 0.39 | 2.27 | 0.26 |
| Bill Depth | 2.59 | 0.27 (0.25) | 2.34 | 0.33 | 2.68 | 0.22 |
| Kipp's distance | 6.99 | 0.04 (0.04) | 6.96 | 0.04 | 7.01 | 0.03 |
| Wing Chord Length | 8.09 | 0.02 (0.02) | 8.10 | 0.02 | 8.06 | 0.02 |
| Tail Length | 0.87 | 0. 62 (0.62) | 0.72 | 0.68 | 0.97 | 0.54 |
| Bill PPC1 | 0.23 | 0.87 (0.86) | 0.23 | 0.88 | 0.23 | 0.87 |
| Bill PPC2 | 8.77 | 0.02 (0.02) | 8.77 | 0.02 | 8.77 | <0.01 |
| CV Bill Length | 3.02 | 0.21 (0.20) | 3.05 | 0.23 | 3.02 | 0.18 |
| CV Bill Width | 3.13 | 0.18 (0.22) | 3.13 | 0.22 | 3.13 | 0.17 |
| CV Bill Depth | 0.60 | 0.71 (0.70) | 0.60 | 0.72 | 0.60 | 0.68 |
| CV Tarsus | 5.88 | 0.06 (0.06) | 5.88 | 0.06 | 5.89 | 0.06 |
| CV Kipp's distance | 0.38 | 0.81 (0.80) | 0.38 | 0.82 | 0.38 | 0.80 |
| CV Wing Chord Length | 0.16 | 0.92 (0.90) | 0.16 | 0.92 | 0.16 | 0.91 |
| CV Tail Length | 0.70 | 0. 67 (0.67) | 0.70 | 0.69 | 0.70 | 0.64 |
| CV Bill PPC1 | <0.01 | 1.0 (1.0) | <0.01 | 1.0 | <0.01 | 1.0 |
| CV Bill PPC2 | 1.02 | 0.54 (0.57) | 1.02 | 0.58 | 1.02 | 0.56 |
