## Supporting Information 2 for "Morphology of migration: Associations between wing, and bill morphology and migration in kingbirds (*Tyrannus*)"

**Supporting Information 2** - ImageJ protocol for measuring long *Tyrannus* tail lengths

We measured tail length as the longest rectrix to the nearest 0.1 cm (Pyle, 1997). For most measurements, we used a Mitutoyo brand IP 67 digital calipers (part number 573-271) with a range of up to 15.24 cm, with 0.00127 cm resolution. For tails longer than 15.24 cm, we used a 30.48 cm stainless steel ruler placed between the two middle rectrices, and when tails were longer than 30.48 cm photos were taken of the tails above a 0.64 x 0.64 cm square grid with the calipers measuring to their extent and ImageJ was used to calculate the full tail length (**Figure S2.1**).

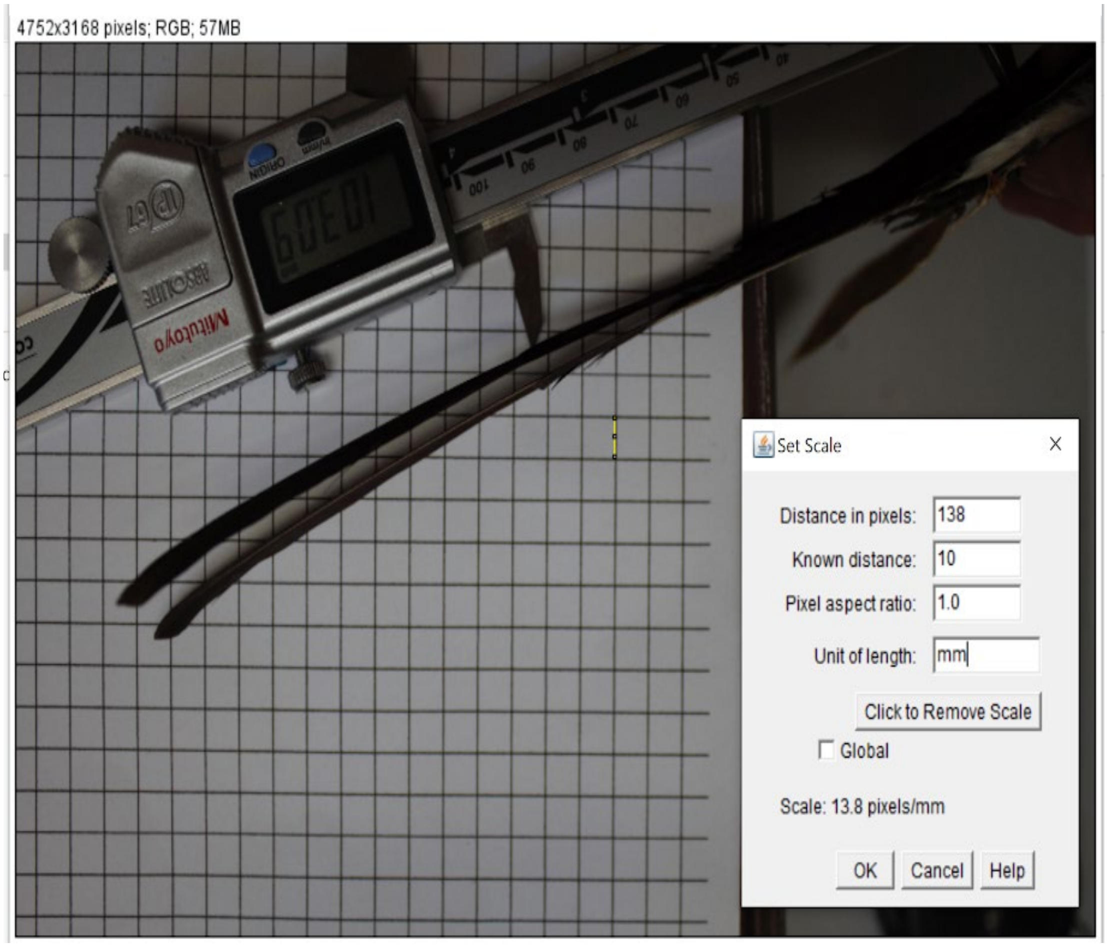


**Figure S2.1**. Image of a *T. savana* tail above 0.64 x 0.64 cm grid. Photos were taken directly above specimens using a RPS brand copy stand (here), Canon Rebel T1i SLR, and Canon 18-55 mm image stabilizer lens. Mitutoyo brand digital calipers were placed between the middle

rectrices to the base of the tail and extended. We prioritized showing the measurement on the digital calipers and the end of the tail for future tail length calculations.

Set the scale using the ‘Set Scale’ tool, we set the scale to 6.4 mm. This is accomplished from the *Straight* tool by clicking and dragging from one corner of a square to another on the grid paper. Then, by clicking Analyze>Set Scale to set the length of the line to 6.4 mm. Finally, click ok.

To measure the tail length, start by clicking from the straight edge of the calipers to the end of the longest tail feather, then click Analyze>Measure (**Figure S3.2**). Sum the length of the measured line (e.g., 147.971 mm) and the caliper reading (e.g., 103.09 mm) and record without decimal places in the data sheet. A ruler was used to measure shorter tails and length estimated to the nearest millimeter. By using two instruments to measure longer tails, we do not assume any increased accuracy and thus estimate to the nearest millimeter as well.

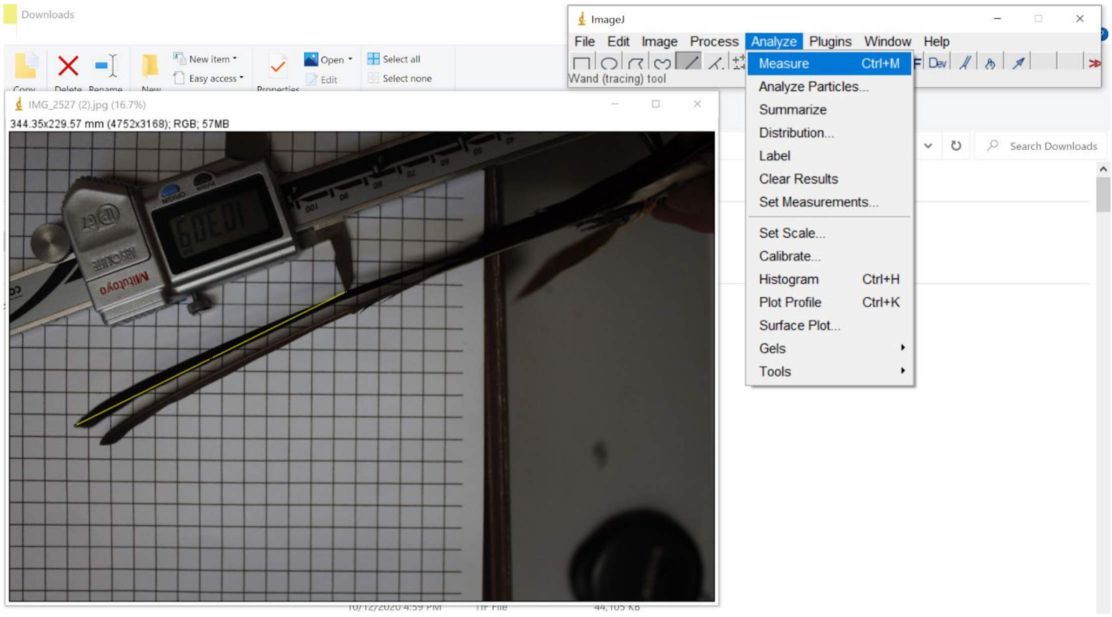

**Figure S2.2**. Measuring the tail length.
