## Supporting Information 3 for "Morphology of migration: Associations between wing, and bill morphology and migration in kingbirds (*Tyrannus*)"

**Supporting Information 3** - Sex-specific results: averages for each morphometric, phylogenetic principal components analysis (PPCA), and phylogenetic ANOVA

This supplement reports the same results as the main text, but for each sex separately. We first report the results for females, followed by the results for males. As the patterns observed were not different than when sexes were lumped we elected to present those results in the main text.

**Table S3.1**. Summary of morphological measurements for **females** from 28 *Tyrannus* OTUs (millimeters).

| OTU | Bill length | Bill width | Bill depth | Tarsus length | Kipp's Index | Wing Cord | Tail Length |
|---|---|---|---|---|---|---|---|
| T. albogularis | 15.03 Â± 0.78 (17) | 8.74 Â± 0.54 (16) | 6.5 Â± 0.35 (16) | 17.68 Â± 0.62 (17) | 26.96 Â± 1.98 (13) | 102.46 Â± 4.28 (13) | 87.06 Â± 3.65 (13) |
| T. caudifasciatus bahamensis | 19.93 Â± 0.46 (4) | 9.24 Â± 0.41 (4) | 7.57 Â± 0.48 (4) | 22.07 Â± 0.5 (4) | 24.21 Â± 2.2 (4) | 105.1 Â± 4.33 (4) | 83.45 Â± 3.2 (4) |
| T. c. caudifasciatus | 18.88 Â± 2.48 (22) | 8.78 Â± 0.9 (22) | 7.3 Â± 0.34 (17) | 22.32 Â± 0.79 (22) | 21.82 Â± 1.7 (19) | 101.31 Â± 2.37 (19) | 83.78 Â± 2.97 (16) |
| T. c. caymanensis | 20.98 Â± 0.52 (2) | 9.22 Â± 0.35 (2) | 7.39 Â± 0.69 (2) | 22.67 Â± 0.66 (3) | 21 Â± 1.48 (3) | 100.46 Â± 0.83 (3) | 85.4 Â± 4.23 (3) |
| T. c. flavescens | 19.84 Â± 0.52 (9) | 8.42 Â± 0.21 (9) | 7.18 Â± 0.14 (9) | 22.26 Â± 0.36 (9) | 22.94 Â± 1.45 (9) | 102.78 Â± 3.27 (9) | 83.44 Â± 2.96 (9) |
| T. c. gabbii | 18.16 Â± 0.63 (13) | 9.02 Â± 0.39 (14) | 6.77 Â± 0.42 (12) | 21.09 Â± 0.63 (14) | 21.33 Â± 1.65 (9) | 102.43 Â± 3.63 (9) | 78.94 Â± 1.59 (8) |
| T. c. jamaicensis | 20.35 Â± 0.9 (4) | 9.1 Â± 0.08 (4) | 7.23 Â± 0.47 (3) | 21.71 Â± 0.36 (4) | 22.43 Â± 2.68 (3) | 100.79 Â± 3.23 (3) | 83.06 Â± 3.29 (2) |
| T. c. taylori | 20.15 Â± 0.85 (26) | 10.07 Â± 0.47 (27) | 8.07 Â± 0.28 (21) | 23.55 Â± 0.67 (27) | 23.47 Â± 1.78 (23) | 111.28 Â± 3.27 (23) | 92.4 Â± 1.97 (21) |
| T. couchii | 16.36 Â± 2.24 (37) | 10.32 Â± 1.04 (36) | 7.78 Â± 0.47 (35) | 19.5 Â± 0.92 (37) | 31.17 Â± 2.07 (25) | 115.53 Â± 4.22 (25) | 92.75 Â± 4.93 (19) |
| T. crassirostris crassirostris | 20.3 Â± 1.12 (11) | 12.56 Â± 0.52 (12) | 10.48 Â± 0.26 (11) | 20.47 Â± 0.61 (13) | 35.51 Â± 3.46 (11) | 125.86 Â± 5.02 (11) | 94.19 Â± 3.3 (9) |
| T. c. pompalis | 20.27 Â± 0.99 (22) | 12.42 Â± 0.87 (23) | 10.68 Â± 0.46 (22) | 20.93 Â± 0.7 (23) | 37.12 Â± 2.81 (18) | 128.42 Â± 3.19 (18) | 94.22 Â± 3.18 (19) |
| T. cubensis | 26.14 Â± 0.69 (17) | 14.33 Â± 0.49 (18) | 11.83 Â± 0.47 (18) | 23.18 Â± 0.88 (18) | 32.88 Â± 2.89 (12) | 127.18 Â± 2.88 (12) | 93.43 Â± 4.67 (6) |
| T. dominicensis dominicensis | 20.58 Â± 1 (37) | 11.12 Â± 0.5 (37) | 8.73 Â± 0.52 (36) | 18.99 Â± 0.54 (41) | 32.02 Â± 2.5 (31) | 111.49 Â± 3.82 (31) | 84.79 Â± 3.14 (24) |
| T. d. vorax | 22.98 Â± 1.01 (28) | 12.67 Â± 0.62 (27) | 9.31 Â± 0.53 (25) | 19.53 Â± 0.78 (32) | 30.73 Â± 1.84 (25) | 113.46 Â± 3.25 (25) | 89.35 Â± 2.61 (16) |
| T. forficatus | 13.28 Â± 0.65 (37) | 7.68 Â± 0.39 (37) | 6.28 Â± 0.3 (37) | 18.97 Â± 0.81 (40) | 38.16 Â± 4.15 (30) | 113.33 Â± 4.69 (30) | 143.74 Â± 23.38 (33) |
| T. melancholicus despotes | 17.26 Â± 1.12 (14) | 9.7 Â± 0.64 (15) | 7.27 Â± 0.58 (11) | 17.61 Â± 1.08 (15) | 27.98 Â± 0.58 (10) | 104.3 Â± 3.2 (10) | 87.14 Â± 2.23 (5) |
| T. m. melancholicus | 17.21 Â± 1.32 (114) | 9.67 Â± 0.6 (114) | 7.49 Â± 0.61 (111) | 18.29 Â± 0.84 (116) | 29.37 Â± 2.31 (88) | 110.41 Â± 2.99 (88) | 89.9 Â± 3.44 (49) |
| T. m. satrapa | 17.58 Â± 1.04 (81) | 10.04 Â± 0.46 (84) | 7.41 Â± 0.38 (83) | 19.32 Â± 0.92 (86) | 27.58 Â± 1.98 (66) | 109.33 Â± 4.12 (66) | 88.45 Â± 3.85 (53) |
| T. niveigularis | 14.63 Â± 0.39 (7) | 8.79 Â± 0.36 (7) | 6.33 Â± 0.35 (7) | 18.35 Â± 0.88 (7) | 24.2 Â± 1.8 (6) | 101.26 Â± 3.44 (6) | 76.71 Â± 2.01 (6) |

| | | | | | | | |
|---|---|---|---|---|---|---|---|
| T. savana circumdatus | 11.48 Â± 0.75<br>(5) | 6.29 Â± 0.37<br>(5) | 5.01 Â± 0.21<br>(5) | 17.49 Â± 0.44<br>(5) | 30.74 Â± 2.08<br>(5) | 100.03 Â± 1.91<br>(5) | 182 Â± 15.62<br>(3) |
| T. s. monachus CA | 11.12 Â± 0.46<br>(19) | 6.66 Â± 0.31<br>(20) | 5.25 Â± 0.27<br>(20) | 17.69 Â± 0.65<br>(20) | 30.13 Â± 2.33<br>(14) | 97.21 Â± 3.64<br>(14) | 175.4 Â± 33.64<br>(8) |
| T. s. monachus SA | 11.35 Â± 0.77<br>(43) | 6.35 Â± 0.45<br>(43) | 5.32 Â± 0.33<br>(40) | 17.24 Â± 0.57<br>(18) | 31.68 Â± 2.42<br>(14) | 100.3 Â± 3.74<br>(33) | 181.42 Â± 36.39<br>(24) |
| T. s. sanctaemartae | 12.03 Â± 0.43<br>(4) | 6.38 Â± 0.24<br>(4) | 5.45 Â± 0.29<br>(4) | 17.69 Â± 0.85<br>(4) | 31.95 Â± 0.78<br>(3) | 99.5 Â± 1.5<br>(3) | 198 Â± NA<br>(1) |
| T. s. savana | 11.03 Â± 0.64<br>(51) | 6.52 Â± 0.53<br>(50) | 5.33 Â± 0.39<br>(46) | 17.71 Â± 1.16<br>(44) | 33.47 Â± 2.01<br>(26) | 100.15 Â± 16.51<br>(33) | 183.51 Â± 31.84<br>(28) |
| T. tyrannus | 13.3 Â± 0.81<br>(36) | 8.09 Â± 0.49<br>(35) | 6.41 Â± 0.37<br>(34) | 18.84 Â± 0.8<br>(36) | 35.5 Â± 2.74<br>(20) | 111.93 Â± 4.28<br>(20) | 78.12 Â± 4.67<br>(17) |
| T. verticalis | 13.14 Â± 0.92<br>(47) | 7.78 Â± 0.46<br>(47) | 6.6 Â± 0.43<br>(45) | 19.53 Â± 0.74<br>(51) | 36 Â± 2.65<br>(42) | 118.56 Â± 4.32<br>(43) | 84.73 Â± 3.06<br>(43) |
| T. vociferans vociferans | 15.18 Â± 0.82<br>(74) | 8.76 Â± 0.43<br>(75) | 7.08 Â± 0.38<br>(73) | 19.83 Â± 0.75<br>(79) | 35.51 Â± 2.75<br>(50) | 124.63 Â± 4.29<br>(50) | 88.22 Â± 2.85<br>(42) |
| T. vociferans xenopterus | 14.64 Â± 0.54<br>(5) | 8.57 Â± 0.2<br>(5) | 6.72 Â± 0.4<br>(5) | 19.65 Â± 0.54<br>(5) | 32.11 Â± 2.71<br>(3) | 120 Â± 1.84<br>(3) | 89.12 Â± 2.22<br>(3) |

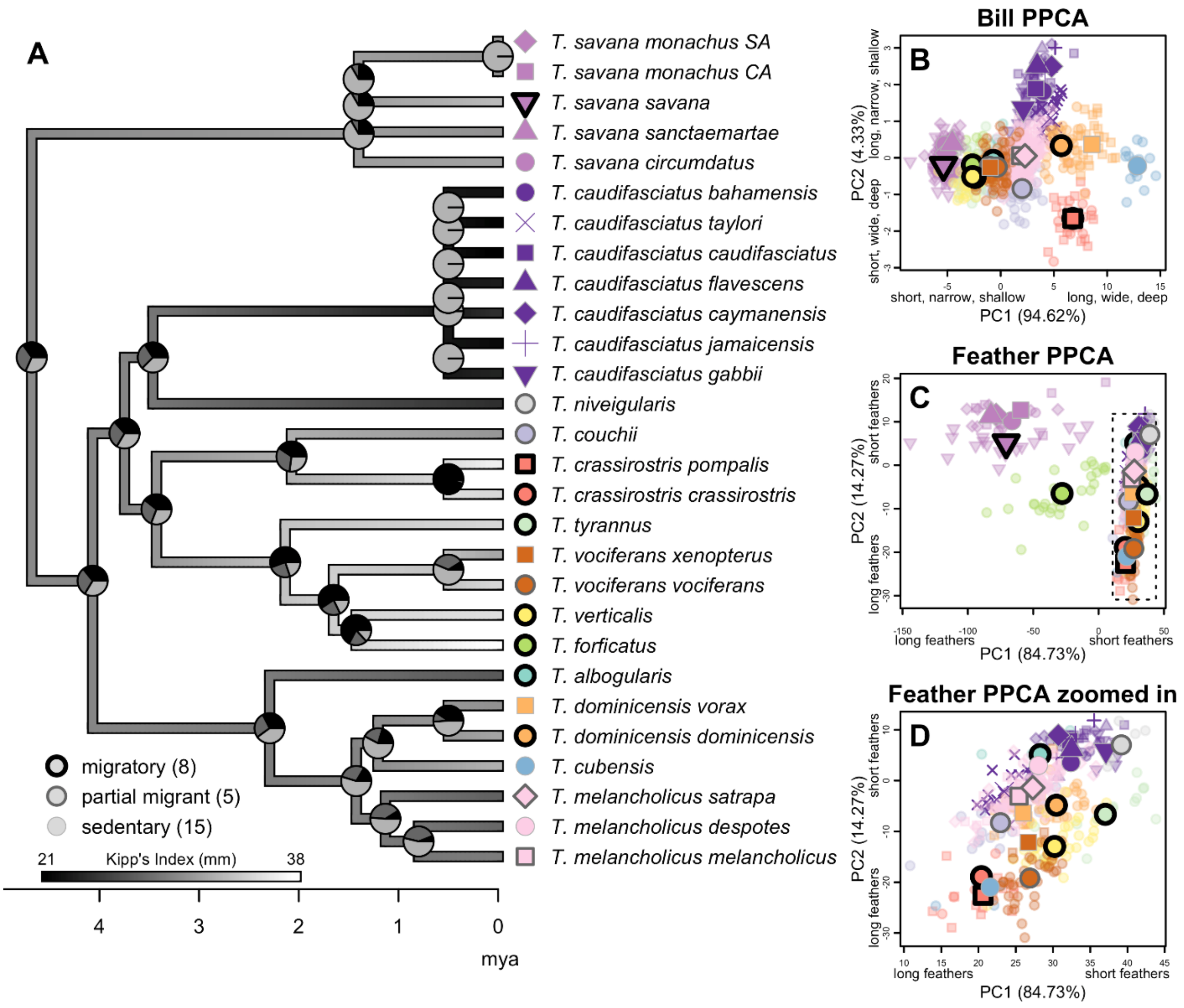


**Figure S3.1**. Ancestral state reconstruction of migration ecology strategy and wing chord length, and ordination of 28 *Tyrannus* OTU adult **females** for bill (length, width, depth) and feather (wing chord length, tail length, and Kipp's distance). A) Kingbird phylogeny showing ancestral state reconstruction of migration ecology strategy (pie charts at each node), and wing chord length (branch greyscale). B-D) PPCA plots showing individuals color-coded to species (identified at the tip labels of the phylogeny), shape-coded to subspecies, and their status as migratory, partially migratory, or sedentary is distinguished via the shape outline. Pie charts for each node in the phylogeny show the ancestral state reconstruction of migration type. E-H) photos of representative kingbird taxa with corresponding color- and shape-coded points in the top right of each photo. Taxa are as follows: E) *Tyrannus savana savana* (photo credit: Rodrigo Conte), F) *Tyrannus caudifasciatus caudifasciatus* (photo credit: Yeray Seminario / Whitehawk), G) *Tyrannus crassirostris pompalis* (photo credit: Martin Molina), and H) *Tyrannus cubensis* (photo credit: Dubio Shapiro).

**Table S3.2**. Results of phylogenetic principal component analysis (PPCA) of bill morphometrics for adult **females**. Percent variance explained by each eigenvector is in brackets for each principal component.

| **Loadings** | **Bill PPC1 (94.13 %)** | **Bill PPC2 (4.72 %)** | **lambda** |
|---|---|---|---|
| Bill length | 0.99 | 0.15 | 0.91 |
| Bill width | 0.93 | -0.33 | |
| Bill depth | 0.94 | -0.28 | |

**Table S3.3**. T and P values from phylogenetic anova analysis for adult **females**. Values shown in brackets are from phylogenetic ANOVA with Bonferroni correction to account for multiple hypothesis testing. No Bonferroni correction was done for the assessment of Tail length because it was analyzed alone with two taxa removed due to missing data (*T. savana sanctaemartae, T. caudfasciatus jamaicensis*). Significant results are in bold.

| **morphometric** | **Migratory v. partially migratory** | | **Migratory v. sedentary** | | **Partially migratory v. sedentary** | |
|---|---|---|---|---|---|---|
| | T | P (adj. p) | T | P (adj. p) | T | P (adj. p) |
| Bill length | 0.33 | 0.74 (1.0) | 0.044 | 0.97 (1.0) | 0.33 | 0.83 (1.0) |
| CV Bill length | 0.34 | 0.71 (1.0) | 1.94 | 0.20 (0.65) | 1.27 | 0. 37 (1.0) |
| Bill width | 0.24 | 0.80 (1.0) | 1.86 | 0.25 (0.75) | 1.84 | 0.20 (0.55) |
| CV Bill width | 1.26 | 0.20 (0.52) | 2.24 | 0.14 (0.49) | 0.51 | 0.72 (1.0) |
| Bill depth | 0.50 | 0.60 (1.0) | 2. 31 | 0.15 (0.42) | 1.41 | 0.32 (0.88) |
| CV Bill depth | 0.79 | 0.42 (1.0) | 0.84 | 0.59 (1.0) | 0.15 | 0.91 (1.0) |
| Bill PC1 (size) | 0.197 | 0.93 (1.0) | 0.55 | 0.74 (1.0) | 0.36 | 0.80 (1.0) |
| CV Bill PC1 (size) | 0.13 | 0.89 (1.0) | 0. 31 | 0.84 (1.0) | 0.12 | 0.94 (1.0) |
| Bill PC2 (shape) | 0.64 | 0.54 (1.0) | 3.82 | **0.024 (0.045)** | 2.53 | 0.074 (0.20) |
| CV Bill PC2 (shape) | 1.31 | 0.18 (0.54) | 3.63 | **0.018** (0.075) | 1.63 | 0.26 (0.74) |
| Kipp’s distance | 1.23 | 0.046 (0.17) | 4 .09 | **0.005 (0.024)** | 1.23 | 0.38 (1.0) |
| CV Kipp’s Index | 0.15 | 0.88 (1.0) | 1.40 | 0.41 (1.0) | 1.02 | 0.46 (1.0) |
| Wing chord length | 0.54 | 0.58 (1.0) | 3. 48 | **0.012 (0.054)** | 2.65 | **0.047** (0.162) |
| CV Wing chord length | 0.21 | 0.84 (1.0) | 2.24 | 0.16 (0.43) | 2.13 | 0.13 (0.36) |
| Tail length | 0.94 | 0.34 (1.0) | 0.44 | 0.77 (1.0) | 1.41 | 0.30 (1.0) |

| | | | | | | |
|---|---|---|---|---|---|---|
| Kipp’s Index | 1.99 | 0.059 (0.16) | 2.99 | 0.060 (0.17) | 0.34 | 0.80 (1.0) |
| CV Kipp’s Index | 0.42 | 0.66 (1.0) | 0.38 | 0.80 (1.0) | 0.14 | 0.92 (1.0) |
| CV Tail length | 1.32 | 0.1 7 | 0.83 | 0.61 | 0.72 | 0.59 |
| CV Tarsus length | 0.98 | 0.31 (0.89) | 1.96 | 0.21 (0.66) | 2.74 | **0.048** (0.13) |

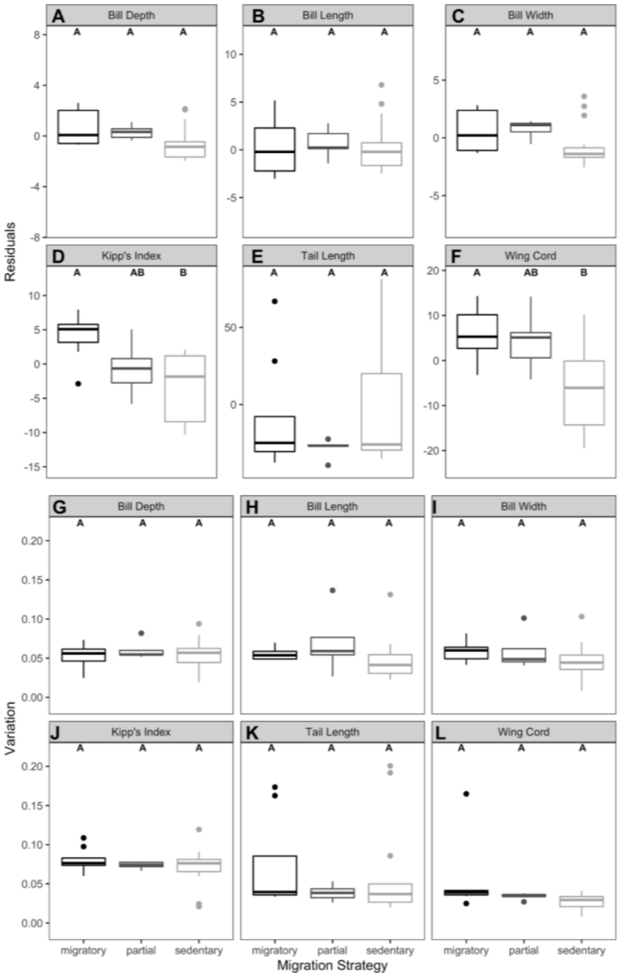


**Figure S3.2**. Phylogenetic ANOVA results comparing residuals of bill and feather morphometrics (A-F), and coefficient of variation in bill and feather morphometrics (G-L) across migratory, partially migratory and sedentary *Tyrannus* OTU adult **females**. Significant differences are shown by different letters: A versus B.

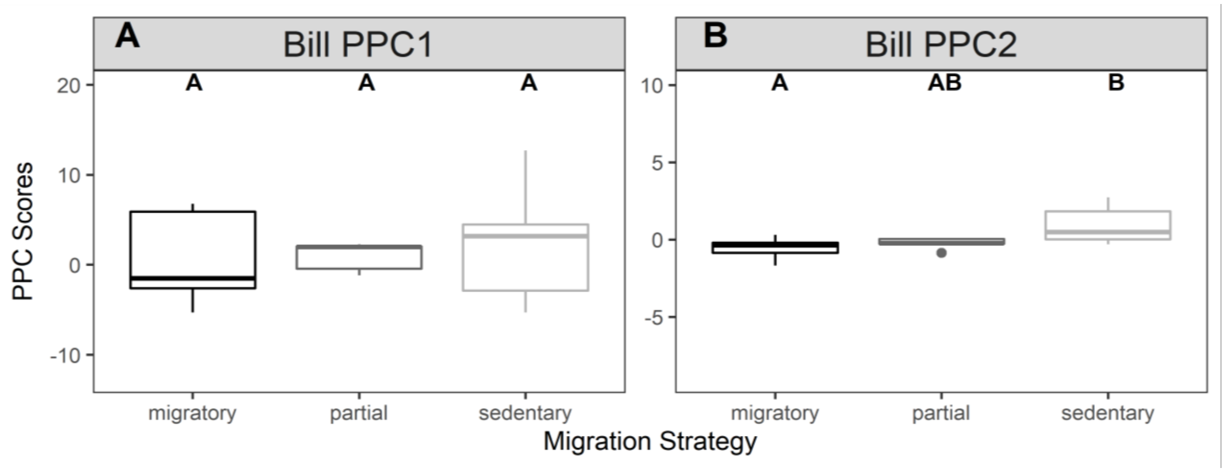


**Figure S3.3**. Phylogenetic ANOVA results comparing bill phylogenetic principal component (PPC) scores across migratory, partially migratory, and sedentary *Tyrannus* OTU adult **females**. Significant differences shown by different letters: A versus B.

---------------------------------------------------------------------------------------------------------------------

**Table S3.4**. Summary of morphological measurements for **males** from 28 *Tyrannus* OTUs (millimeters).

| OTU | Bill length | Bill width | Bill depth | Tarsus length | Kipp's Index | Wing Cord | Tail Length |
|---|---|---|---|---|---|---|---|
| T. albogularis | 15.24 Â± 0.7 (19) | 8.51 Â± 0.4 (19) | 6.24 Â± 0.36 (17) | 17.48 Â± 0.86 (19) | 30.08 Â± 1.32 (13) | 107.72 Â± 3.99 (13) | 94.98 Â± 4.75 (13) |
| T. couchii | 20.24 Â± 1.04 (7) | 9.47 Â± 0.44 (7) | 7.59 Â± 0.33 (7) | 22.45 Â± 0.3 (7) | 25.98 Â± 1.25 (6) | 109.28 Â± 2.48 (6) | 85.69 Â± 2.59 (6) |
| T. cubensis | 19.41 Â± 1.84 (47) | 8.87 Â± 0.75 (47) | 7.21 Â± 0.53 (39) | 22.58 Â± 0.67 (48) | 24.21 Â± 2.06 (35) | 104.94 Â± 2.1 (36) | 86.19 Â± 2.43 (31) |
| T. forficatus | 21.76 Â± 0.96 (10) | 9.46 Â± 0.23 (9) | 7.42 Â± 0.21 (9) | 22.58 Â± 0.64 (12) | 22.11 Â± 1.96 (8) | 104.14 Â± 2.56 (8) | 89.05 Â± 3.28 (7) |
| T. niveigularis | 20.03 Â± 0.78 (12) | 8.51 Â± 0.22 (12) | 7.18 Â± 0.3 (12) | 22.88 Â± 0.36 (12) | 23.94 Â± 0.68 (12) | 106.21 Â± 2.41 (12) | 84.25 Â± 1.96 (12) |
| T. tyrannus | 18.39 Â± 0.67 (17) | 8.79 Â± 0.41 (17) | 6.66 Â± 0.31 (16) | 20.79 Â± 0.74 (17) | 23.22 Â± 1.14 (14) | 103.46 Â± 3.64 (15) | 81.72 Â± 2.53 (14) |
| T. verticalis | 19.89 Â± 1.58 (7) | 9.11 Â± 0.42 (7) | 7.03 Â± 0.54 (7) | 21.7 Â± 0.53 (7) | 22.46 Â± 1.68 (6) | 99.82 Â± 3.3 (6) | 84.42 Â± 4.23 (6) |
| T. caudifasciatus bahamensis | 20.49 Â± 0.86 (40) | 9.96 Â± 0.57 (40) | 7.83 Â± 0.34 (30) | 23.74 Â± 0.74 (40) | 24.91 Â± 1.46 (36) | 113.54 Â± 3.24 (36) | 92.51 Â± 2.46 (35) |
| T. c. caudifasciatus | 16.54 Â± 1.42 (49) | 10.08 Â± 0.66 (48) | 7.52 Â± 0.42 (44) | 19.45 Â± 0.62 (51) | 31.17 Â± 2.89 (34) | 116.75 Â± 5.86 (34) | 94.85 Â± 5.24 (30) |
| T. c. caymanensis | 19.28 Â± 0.98 (9) | 12.31 Â± 0.6 (8) | 10.07 Â± 0.45 (8) | 20.2 Â± 0.87 (9) | 37.08 Â± 3.5 (7) | 128.5 Â± 5.76 (7) | 98.69 Â± 3.75 (4) |
| T. c. flavescens | 20.28 Â± 0.82 (27) | 12.24 Â± 0.58 (27) | 10.46 Â± 0.49 (26) | 20.67 Â± 0.81 (27) | 38.07 Â± 3.71 (22) | 130.22 Â± 3.74 (22) | 95.81 Â± 3.61 (24) |
| T. c. gabbii | 26.08 Â± 0.82 (14) | 13.92 Â± 0.38 (13) | 11.55 Â± 0.47 (14) | 22.86 Â± 0.6 (14) | 34.45 Â± 1.85 (11) | 131.01 Â± 3.45 (11) | 97.85 Â± 5.23 (10) |
| T. c. jamaicensis | 20.55 Â± 1.13 (57) | 10.94 Â± 0.54 (53) | 8.55 Â± 0.53 (49) | 18.61 Â± 0.62 (59) | 33.91 Â± 2.48 (39) | 113.62 Â± 3.62 (38) | 87.24 Â± 3.45 (31) |

| | | | | | | | |
|---|---|---|---|---|---|---|---|
| T. c. taylori | 22.68 Â± 1.33 (33) | 12.6 Â± 0.7 (32) | 9.34 Â± 0.64 (29) | 19.21 Â± 0.6 (35) | 31.8 Â± 2.77 (23) | 115.09 Â± 4.26 (23) | 89.68 Â± 4.55 (16) |
| T. crassirostris crassirostris | 13.65 Â± 0.54 (49) | 7.61 Â± 0.34 (51) | 6.19 Â± 0.34 (48) | 19.14 Â± 0.7 (55) | 45.22 Â± 5.22 (47) | 121.46 Â± 4.99 (47) | 206.17 Â± 43.48 (52) |
| T. c. pompalis | 17.66 Â± 0.71 (35) | 9.69 Â± 0.49 (35) | 7.54 Â± 0.5 (30) | 17.83 Â± 0.93 (36) | 32.37 Â± 2.51 (22) | 109.84 Â± 11.26 (22) | 93.58 Â± 2.95 (11) |
| T. dominicensis dominicensis | 17.52 Â± 1.32 (134) | 9.71 Â± 0.48 (132) | 7.44 Â± 0.6 (128) | 18.09 Â± 0.7 (134) | 31.34 Â± 2.6 (103) | 114.11 Â± 3.88 (104) | 93.28 Â± 5.07 (74) |
| T. d. vorax | 17.65 Â± 1.08 (107) | 9.87 Â± 0.64 (107) | 7.27 Â± 0.38 (105) | 19.09 Â± 0.89 (110) | 29.39 Â± 2.18 (69) | 112.73 Â± 3.81 (69) | 92.88 Â± 4.44 (56) |
| T. melancholicus despotes | 14.68 Â± 0.62 (14) | 8.51 Â± 0.54 (15) | 6.05 Â± 0.3 (15) | 18.16 Â± 0.77 (15) | 25.22 Â± 2.1 (14) | 101.72 Â± 3.76 (14) | 77.28 Â± 2.54 (15) |
| T. m. melancholicus | 12.31 Â± 0.46 (19) | 6.35 Â± 0.26 (19) | 5.23 Â± 0.26 (19) | 17.61 Â± 0.34 (19) | 34.72 Â± 3.34 (17) | 105.26 Â± 3.98 (17) | 229.75 Â± 21.9 (16) |
| T. m. satrapa | 11.28 Â± 0.49 (36) | 6.39 Â± 0.3 (33) | 5.2 Â± 0.36 (32) | 17.28 Â± 0.69 (36) | 34.01 Â± 1.89 (17) | 104.22 Â± 3.48 (17) | 270.67 Â± 27.31 (12) |
| T. savana circumdatus | 11.75 Â± 0.62 (47) | 6.34 Â± 0.4 (46) | 5.35 Â± 0.32 (40) | 17.43 Â± 0.73 (24) | 34.27 Â± 2.23 (20) | 100.79 Â± 17.49 (38) | 247.54 Â± 44.86 (31) |
| T. s. monachus CA | 12.04 Â± 0.26 (9) | 6.08 Â± 0.19 (9) | 5.29 Â± 0.2 (9) | 17.58 Â± 0.44 (9) | 36.39 Â± 3.19 (5) | 105.5 Â± 4.58 (5) | 239 Â± 48.94 (8) |
| T. s. monachus SA | 11.15 Â± 0.68 (76) | 6.48 Â± 0.51 (79) | 5.26 Â± 0.44 (68) | 17.76 Â± 0.88 (59) | 36.73 Â± 4.17 (37) | 108.06 Â± 6.16 (53) | 216.71 Â± 52.54 (44) |
| T. s. sanctaemartae | 13.8 Â± 0.63 (61) | 8.12 Â± 0.94 (61) | 6.38 Â± 0.43 (60) | 18.75 Â± 0.67 (61) | 38.27 Â± 2.73 (44) | 115.78 Â± 4.85 (45) | 79.22 Â± 3.92 (38) |
| T. s. savana | 13.38 Â± 0.96 (50) | 7.68 Â± 0.43 (50) | 6.45 Â± 0.47 (49) | 19.43 Â± 0.79 (54) | 41.34 Â± 3.4 (52) | 125.98 Â± 4.3 (52) | 90.04 Â± 3.91 (50) |
| T. vociferans vociferans | 14.95 Â± 1.17 (96) | 8.54 Â± 0.56 (96) | 6.88 Â± 0.54 (91) | 19.76 Â± 0.69 (108) | 38.88 Â± 3 (62) | 128.58 Â± 13.62 (63) | 90.5 Â± 3.62 (56) |
| T. v. xenopterus | 14.8 Â± 0.88 (7) | 8.21 Â± 0.48 (7) | 6.56 Â± 0.34 (7) | 19.91 Â± 0.67 (7) | 39.87 Â± 4.54 (3) | 130.19 Â± 6.96 (3) | 91.77 Â± 3.75 (4) |

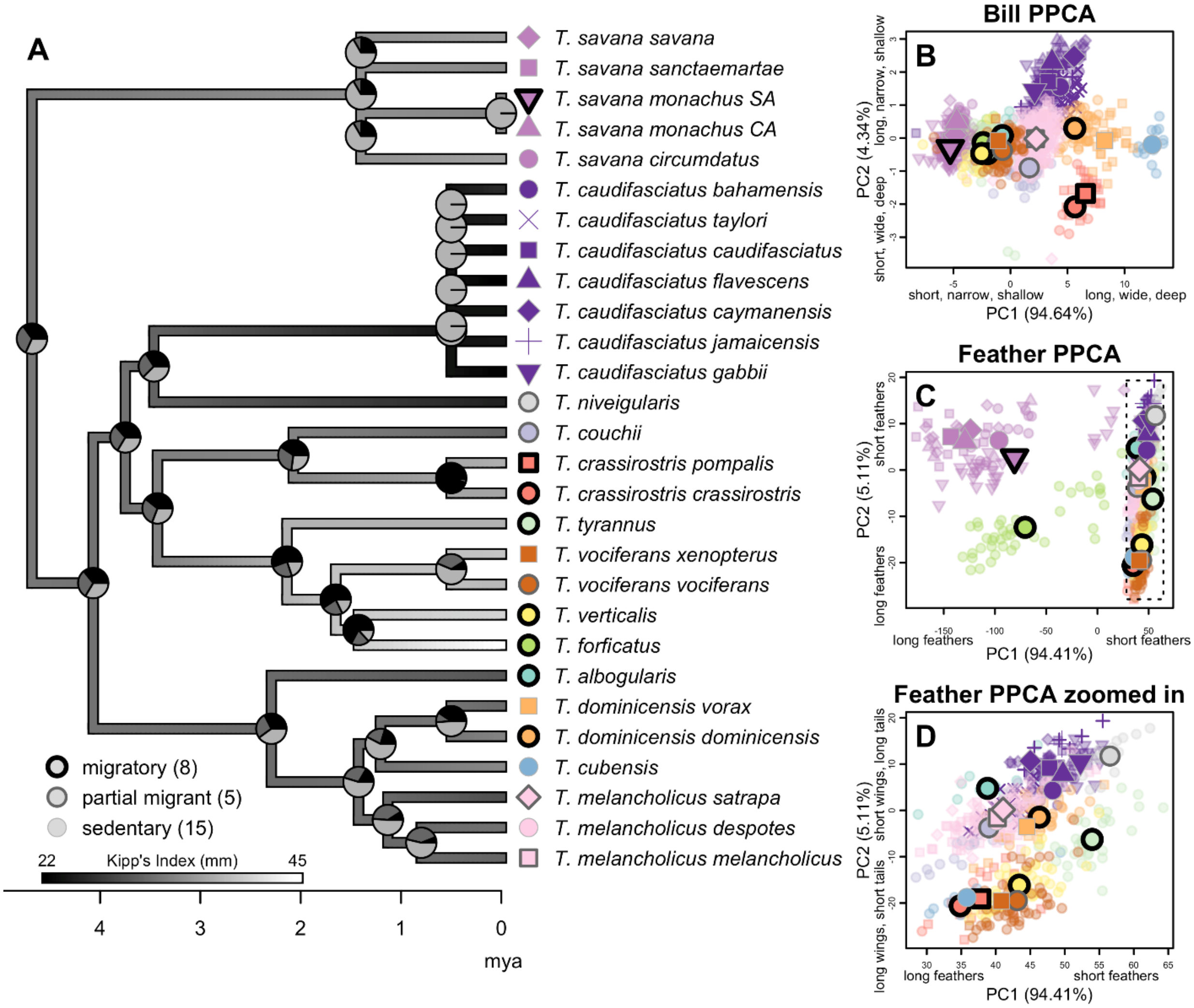


**Figure S3.4**. Ancestral state reconstruction of migration strategy and wing chord length, and ordination of 28 *Tyrannus* OTU adult **males** for bill (length, width, depth) and feather (wing chord length and Kipp's distance). A) Tyrannus phylogeny showing ancestral state reconstruction of migration strategy (pie charts at each node), and wing chord length (branch greyscale). B-D) PPCA plots showing individuals color-coded to species (identified at the tip labels of the phylogeny), shape-coded to subspecies, and their status as migratory, partially migratory or sedentary is distinguished via the shape outline. Pie charts for each node in the phylogeny show the ancestral state reconstruction of migration type. E-H) photos of representative kingbird taxa with corresponding color- and shape-coded points in the top right of each photo. Taxa are as follows: E) *Tyrannus savana savana* (photo credit: Rodrigo Conte), F) *Tyrannus caudifasciatus caudifasciatus* (photo credit: Yeray Seminario / Whitehawk), G) *Tyrannus crassirostris pompalis* (photo credit: Martin Molina), and H) *Tyrannus cubensis* (photo credit: Dubio Shapiro).

**Table S3.5**. Results of phylogenetic principal component analysis (PPCA) of bill morphometrics for adult **males**. Percent variance explained by each eigenvector is in brackets for each principal component.

| **Loadings** | **Bill PPC1 (94.83%)** | **Bill PPC2 (4.24%)** | **lambda** |
|---|---|---|---|
| Bill length | 0.99 | 0.14 | 0.97 |
| Bill width | 0.93 | -0.33 | |
| Bill depth | 0.94 | -0.26 | |

**Table S3.6**. T and P values from phylogenetic ANOVA for **males**. Significant results are in bold.

| **morphometric** | **Migratory v. partially migratory** | | **Migratory v. sedentary** | | **Partially migratory v. sedentary** | |
|---|---|---|---|---|---|---|
| | T | P | T | P | T | P |
| Bill length | 0.31 | 0.76 (1.0) | 0.10 | 0.95 (1.0) | 0.26 | 0.86 (1.0) |
| CV Bill length | 1.29 | 0.20 (0.55) | 0.95 | 0.55 (1.0) | 2.23 | 0.11 (0.38) |
| Bill width | 0.17 | 0.85 (1.0) | 1.47 | 0.33 (1.0) | 1.43 | 0.29 (0.86) |
| CV Bill width | 0.67 | 0.46 (1.0) | 2.08 | 0.16 (0.52) | 1.02 | 0.49 (1.0) |
| Bill depth | 0.48 | 0.63 (1.0) | 2.02 | 0.20 (0.65) | 1.18 | 0.39 (1.0) |
| CV Bill depth | 0.22 | 0.83 (1.0) | 0.72 | 0.65 (1.0) | 0.84 | 0.57 (1.0) |
| Bill PC1 (size) | 0.072 | 0.94 (1.0) | 0.63 | 0.71 (1.0) | 0.45 | 0.75 (1.0) |
| CV Bill PC1 (size) | 0.028 | 0.98 (1.0) | 0.20 | 0.88 (1.0) | 0.20 | 0.88 (1.0) |
| Bill PC2 (shape) | 0.63 | 0.53 (1.0) | 3.89 | **0.006 (0.018)** | 2.60 | **0.047 (0.16)** |
| CV Bill PC2 (shape) | 0.11 | 0.89 (1.0) | 1.10 | 0.50 (1.0) | 0.81 | 0.57 (1.0) |
| Kipp's distance | 2.08 | 0.039 (0.099) | 3.51 | **0.019 (0.06)** | 0.68 | 0.63 (1.0) |
| CV Kipp's distance | 0.14 | 0.88 (1.0) | 0.72 | 0.66 (1.0) | 0.76 | 0.60 (1.0) |
| Wing chord length | 0.79 | 0.43 (1.0) | 3.61 | **0.035 (0.057)** | 2.18 | 0.12 (0.32) |
| CV Wing chord length | 0.84 | 0.38 (1.0) | 0.46 | 0.76 (1.0) | 0.54 | 0.72 (1.0) |
| Tail length | 0.93 | 0.30 (0.95) | 0.30 | 0.85 (1.0) | 1.34 | 0.34 (1.0) |
| CV Tail length | 1.00 | 0.30 (0.94) | 0.50 | 0.77 (1.0) | 0.69 | 0.64 (1.0) |
| Kipp's Index | 1.84 | 0.055 (0.22) | 2.58 | 0.10 (0.25) | 0.15 | 0.91 (1.0) |
| CV Kipp's Index | 1.84 | 0.066 (0.21) | 0.12 | 0.93 (1.0) | 1.93 | 0.17 (0.53) |
| CV Tarsus length | 0.40 | 0.70 (1.0) | 3.13 | **0.054** (0.14) | 2.21 | 0.11 (0.34) |

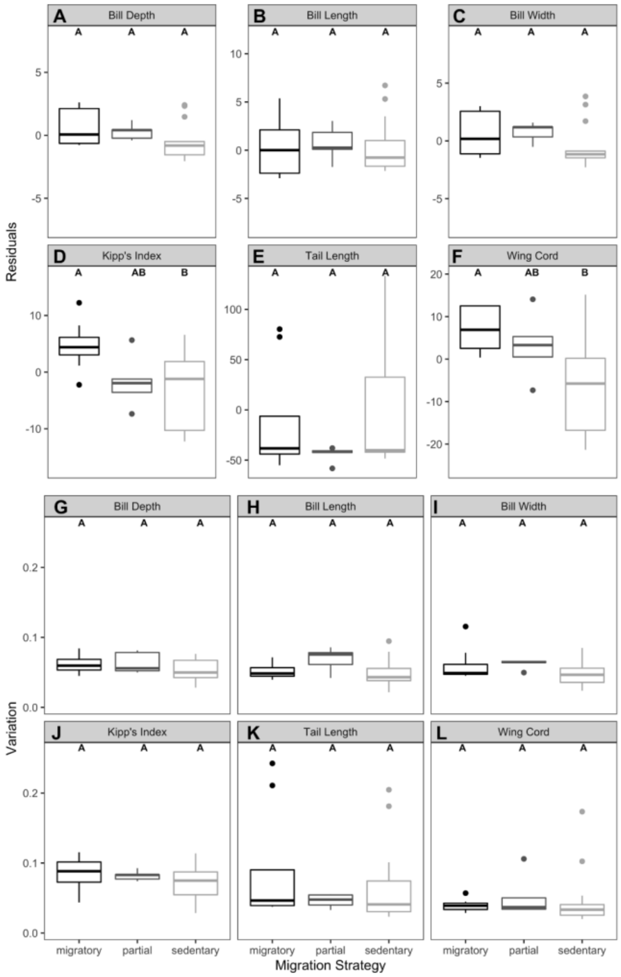


**Figure S3.5**. Phylogenetic ANOVA results comparing residuals of bill and feather morphometrics (A-F), and coefficient of variation in bill and feather morphometrics (G-L) across migratory, partially migratory and sedentary *Tyrannus* OTU adult **males**. Significant differences are shown by different letters: A versus B.

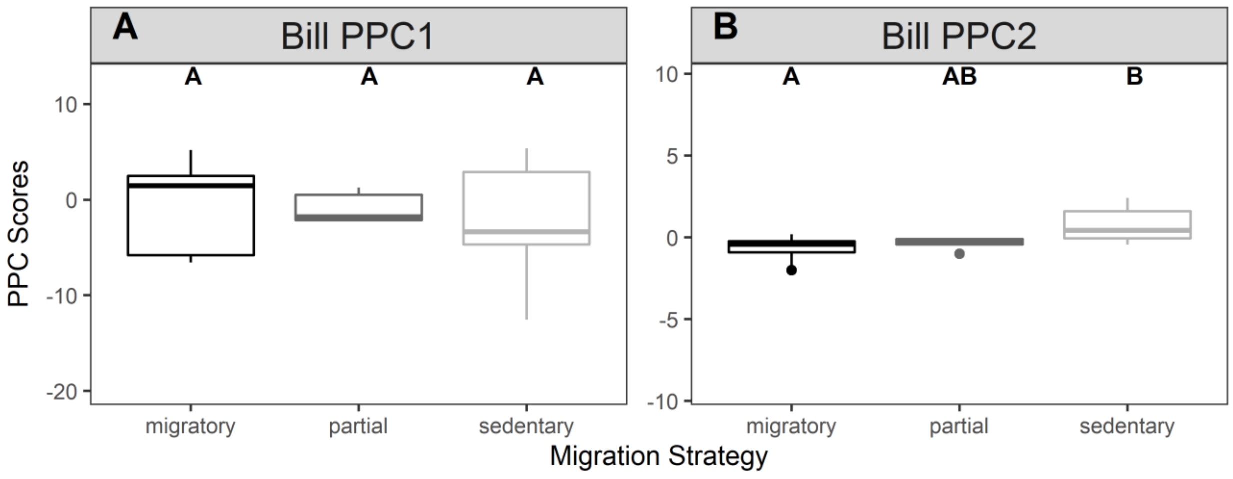


**Figure S3.6**. Phylogenetic ANOVA results comparing bill phylogenetic principal component (PPC) scores across migratory, partially migratory, and sedentary *Tyrannus* OTU adult **males**. Significant differences shown by different letters: A versus B.
